# Correlative SMLM and electron tomography reveals endosome nanoscale domains

**DOI:** 10.1101/629147

**Authors:** Christian Franke, Urska Repnik, Sandra Segeletz, Nicolas Brouilly, Yannis Kalaidzidis, Jean-Marc Verbavatz, Marino Zerial

**Author notes:** authors contributed equally (Christian Franke, Urska Repnik, Sandra Segeletz, Nicolas Brouilly). Corresponding author: Marino Zerial; Max Planck Institute of Molecular Cell Biology and Genetics, MPI-CBG, Pfotenhauerstrasse 108, 01307 Dresden, Germany.

## Abstract

Many cellular organelles, including endosomes, show compartmentalization into distinct functional domains, which however cannot be resolved by diffraction-limited light microscopy. Single molecule localization microscopy (SMLM) offers nanoscale resolution but data interpretation is often inconclusive when the ultrastructural context is missing. Correlative light electron microscopy (CLEM) combining SMLM with electron microscopy (EM) enables correlation of functional sub-domains of organelles in relation to their underlying ultrastructure at nanometer resolution. However, the specific demands for EM sample preparation and the requirements for fluorescent single-molecule photo-switching are opposed. Here, we developed a novel superCLEM workflow that combines triple-colour SMLM (*d*STORM & PALM) and electron tomography using semi-thin Tokuyasu thawed cryosections. We applied the superCLEM approach to directly visualize nanoscale compartmentalization of endosomes in HeLa cells. Internalized, fluorescently labelled Transferrin and EGF were resolved into morphologically distinct domains within the same endosome. We found that the small GTPase Rab5 is organized in nano-domains on the globular part of early endosomes. The simultaneous visualization of several proteins in functionally distinct endosomal sub-compartments demonstrates the potential of superCLEM to link the ultrastructure of organelles with their molecular organization at nanoscale resolution.

**Synopsis:** Suborganelle compartmentalization cannot be resolved by diffraction limited light microscopy and not interpreted without knowledge of the underlying ultrastructure. This work shows a novel superCLEM workflow that combines multi-colour single-molecule localization-microscopy with electron tomography to map several functional domains on early endosomes. superCLEM reveals that the small GTPase Rab5 is organized in nano-domains largely devoid from cargo molecules Transferrin and EGF and opens new possibilities to perform structure-function analysis of organelles at the nanoscale.

## Introduction

Cellular organelles have a characteristic size, shape and a morphologically distinguishable sub-domain organization^1–6^. Such sub-domains are distinct in protein and lipid composition and provide the microenvironment necessary for coordinated biochemical reactions. For example, early endosomes partition cargo molecules with different intracellular fates into distinct sub-domains. Recycling cargo such as Transferrin (Tfn) is sorted into tubules, which pinch off from the endosome and fuse with the plasma membrane. Cargo to be degraded, such as the epidermal growth factor (EGF) and the low-density-lipoprotein (LDL), accumulates in the globular portion of the endosomes^7,8^. EGF remains bound to its receptor and is sorted into intraluminal vesicles (ILVs), whereas LDL is released from its receptor into the lumen of the endosome. The past thirty years have seen growing progress towards the identification of the molecules underlying the structural and functional properties of membrane organelles and their communication^9–13^. Their position in relation to the ultrastructure of the organelle is less understood but of particular significance in order to relate molecular machineries to their nano-environment. For example, confocal microscopy has provided some evidence that Rab GTPases, key regulators of organelle biogenesis and trafficking^14–16^, are compartmentalized into morphologically-distinct domains within early and late endosomes^17,18^. Specifically, the function of Rab5 in regulation of early endosome tethering and fusion^19,20^ is thought to depend on its concentration within sub-endosomal domains^15^. However, a conclusive proof of the existence of such Rab-domains, especially in relation to other functional domains (e.g. cargo sorting), within the same endosome is lacking, mainly because of the diffraction-limited resolution of confocal microscopy.

Electron microscopy (EM) offers nanoscale resolution, but only a few methods are available for localizing specific molecules by EM, and even these are limited by significant constraints. Immuno-gold based protocols generally yield low labelling efficiency because only antigens exposed on the surface of a section are accessible to antibodies and the binding of gold-conjugated probes is hampered by steric hindrance and electron repulsion^21,22^. Some cargo, such as Tfn, may be coupled to colloidal gold particles, which can then be internalized by cells prior to fixation. However, association of a ligand with a gold particle may reduce its internalization efficiency due to steric hindrance. Moreover, coupling to gold may alter the normal intracellular pathway of ligands, as in the case of the Tfn-gold probe, which was observed not to recycle^23^. Therefore, colloidal gold-conjugated probes often results in sparse signal in EM, making it difficult to draw definite conclusions on the sub-compartmental organization of organelles. Alternatively, protein tags that can be visualized by EM are available, but can be applied to only one protein at a time and lead to a diffuse signal with low signal/noise ratio (HRP-tagging^24^; MiniSOG-tagging^25^; Apex-tagging^26–28^)

On the other hand, fluorescence labelling approaches are versatile and range from standard immuno-labelling, tagging proteins with fluorescent tags, to small organic dye-conjugated probes. Despite the generally high labelling efficiency, the diffraction limited resolution of classical light microscopy (≥200 nm) does not allow the visualization of structural features at the nanoscale. Fluorescence based <u>s</u>ingle-<u>m</u>olecule localization-<u>m</u>icroscopy (SMLM) methods, such as <u>p</u>hoto-<u>a</u>ctivation localization <u>m</u>icroscopy (PALM)^30^ and *<u>d</u>*irect <u>s</u>tochastic <u>o</u>ptical reconstruction <u>m</u>icroscopy (*d*STORM)^31^, provide spatial resolutions well below the diffraction barrier and attain structural insight towards the molecular range^32^.

However, despite the improved resolution of SMLM over conventional light microscopy, it is often necessary to visualize the organelle ultrastructure to draw conclusions. Correlative light electron microscopy (CLEM) approaches deploying SMLM can be used to investigate the localization of proteins at the nanoscale in the context of the ultrastructural reference space^33–36^. Yet, single-molecule based CLEM approaches have mostly been limited to one high-resolved colour, either a fluorescent protein or an organic dye. This is largely due to the difficulty of combining multiple photo-switchable markers in a single CLEM workflow with SMLM imaging constraints^29,30,37–39^. Although some dual-colour SMLM based CLEM approaches have been reported^40,41^, they have never been applied to structurally frail structures like endosomes, which are susceptible to ultrastructural damage.

Here, we addressed this challenge by developing a correlative triple-colour SMLM and electron tomography superCLEM workflow that relies on the combination of PALM and *d*STORM with electron tomography for high-resolution three-dimensional ultrastructural analysis, on semi-thin Tokuyasu thawed cryosections. Our results show for the first time the compartmentalization of EGF, Tfn and Rab5 (or LDL) simultaneously in relation to the three-dimensional organelle ultrastructure with nanometer resolution in both imaging modes, demonstrating the applicability of our superCLEM workflow to complex biological questions at the sub-organelle level.

## Results

### Characterization of the experimental cell model

Previous studies provided evidence that Rab5 and other Rab proteins, are compartmentalized on endosomal membranes^15,18^. We therefore aimed to apply triple-colour SMLM to further investigate its sub-endosomal localization relative to Tfn and EGF, as cargo markers of the recycling and degrading routes, respectively. Because membrane permeabilization required for immuno-labelling adversely affects the endosomal ultrastructure, we used Tfn and EGF conjugated to Alexa dyes (Tfn-AF568, EGF-AF647) and genetically tagged Rab5c with the monomeric, reversibly photo-switchable fluorescent protein Dronpa. In comparison to more commonly used photo-switchable fluorescent proteins, Dronpa occupies only one spectral channel, thus facilitating triple colour detection on our setup (see methods). To express Dronpa-Rab5c close to endogenous levels, we generated a HeLa BAC cell line^42^. Three distinct populations of cells were sorted based on their levels of expression, i.e. “low”, “mid” and “high”. The percentage of tagged-to-endogenous Rab5c protein was assessed by Western blotting (Figure 1 A), and Dronpa-Rab5c was approximately 15, 27 and 38% of the total amount of Rab5c protein, respectively.

**Figure 1:**
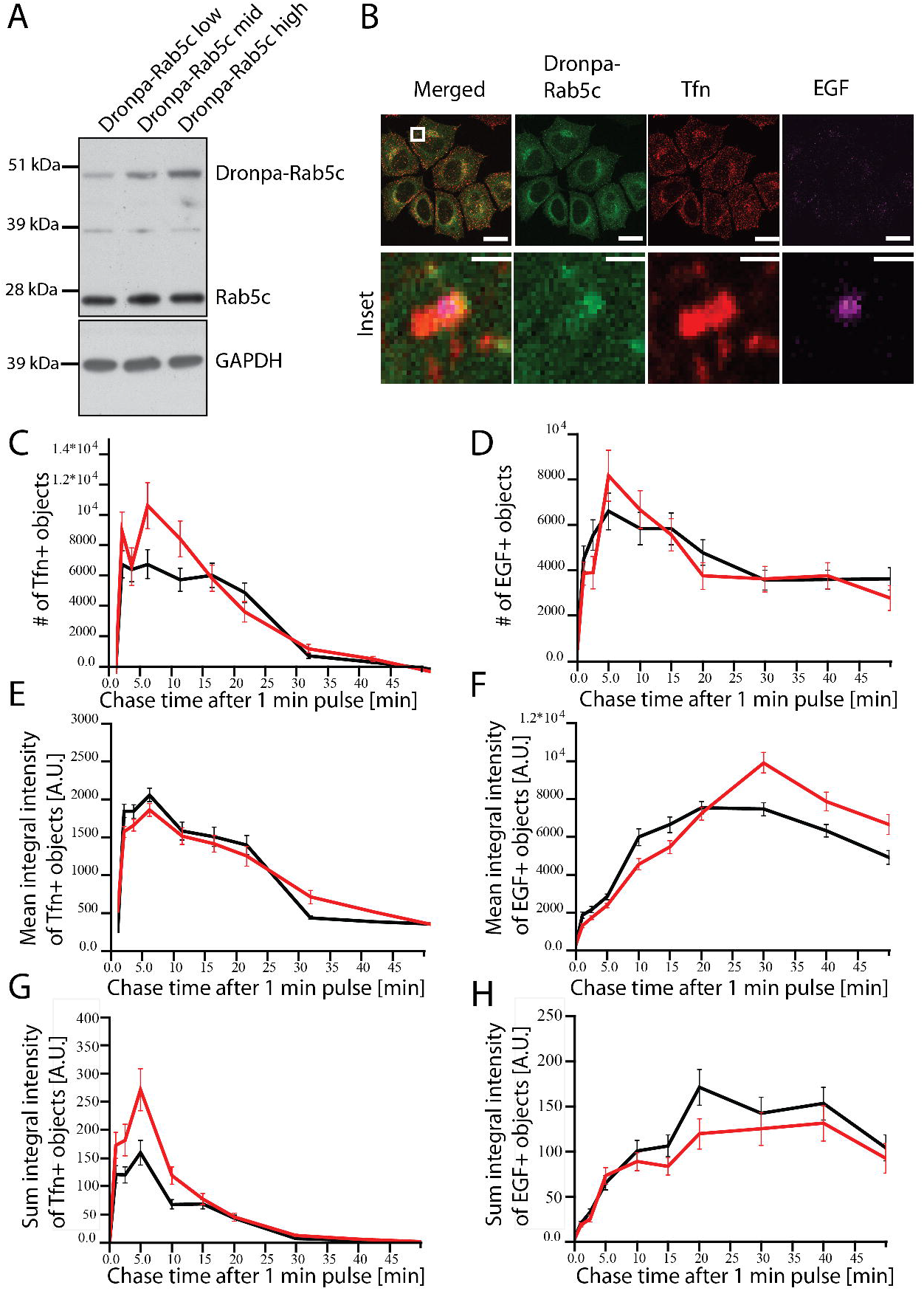
Validation of the Dronpa-Rab5c BAC HeLa cells. (A) Western blot of Dronpa-Rab5c expressing cells sorted by FACS into *low*, *middle* and *high* expression populations. The molecular weights of Dronpa-Rab5c and endogenous Rab5c were confirmed to ~49 kDa and 23 kDa respectively. The percentage of each population was determined to 15 (low), 27 (middle) and 38% (high) relative to the total Rab5c amount. GAPDH was used as a loading control. (B) Representative confocal images of *high*-expression Dronpa-Rab5c cells labelled with a Rab5c antibody. Cells were fed with Tfn and EGF continuously for 15 min prior to fixation. The inlay highlights the indicated boxed area with higher magnification, suggesting a differential spatial distribution of the three proteins. Scale bars: 20 μm, Inlay: 1 μm (C-H) Trafficking kinetics of Tfn (C, E and G) and EGF (D, F and H) for the high expressing Dronpa-Rab5c BAC cell line (Mean±SEM) (C-D) Number of Tfn- (C) and EGF-positive (D) endosomes per masked area. (E-F) Mean integral intensity of Tfn- (E) and EGF-positive (F) endosomes per masked area. (G-H) Sum integral (overall) intensity of Tfn- (G) and EGF-positive endosomes (H) per masked area.

Tagging of proteins with fluorescent labels often causes alterations in endosomal function^43^. To exclude them, we first quantified the trafficking kinetics of Tfn and EGF during a pulse-chase experiment. The uptake of these cargo molecules was measured based on the number and fluorescence intensity of vesicles using quantitative multi-parametric image analysis (QMPIA), as previously reported^43,44^. The “high” population of Dronpa-Rab5c BAC cells showed no substantial alterations in Tfn and EGF trafficking when compared to control HeLa cells and was chosen as a reasonable approximation of endogenous Rab5c levels (Figure 1 C-H). Dronpa-Rab5c was similarly distributed throughout the cell as endogenous Rab5.

In cells incubated with Tfn-AF568 and EGF-AF647 for 15 min, we observed triple-positive EGF, Tfn and Dronpa-Rab5c early endosomes, particularly in the perinuclear region. Confocal microscopy analysis of these regions suggests a differential distribution of Dronpa-Rab5c and the cargo molecules on individual endosomes, which is consistent with the previously reported spatial segregation of different Rab proteins relative to cargo (^17^, Figure 1 B). In addition to the Dronpa-Rab5c cell line, we used a GFP-Rab5c cell line as control throughout our experiments. The endosomal distribution of GFP-Rab5c and the uptake of cargo in the GFP-Rab5c cell line were similar to the Dronpa-Rab5c cells (Suppl. Figure S1).

### Triple colour SMLM resolves endosomal compartmentalization of EGF, Tfn & Rab5

In order to elucidate the compartmentalization of Rab5c in relation to Tfn and EGF, we performed triple colour SMLM on HeLa cells grown on cover slips, after EGF and Tfn uptake for 15 min (Figure 2, Suppl. Figure 2). Similar to confocal imaging, we observed endosomes where all three markers appeared within a certain degree of overlap. EGF was mainly concentrated to a small part of the apparent endosomal area often exhibiting a granular structure. Further, small EGF hotspots were visible at the centre of the structures. On these endosomes, Tfn covered a larger area than EGF but was mainly segregated from it in prominent elongated structures, ranging from short stubs (Figure 2 A) to several 100 nm in length (Figure 2 C,D).

**Figure 2:**
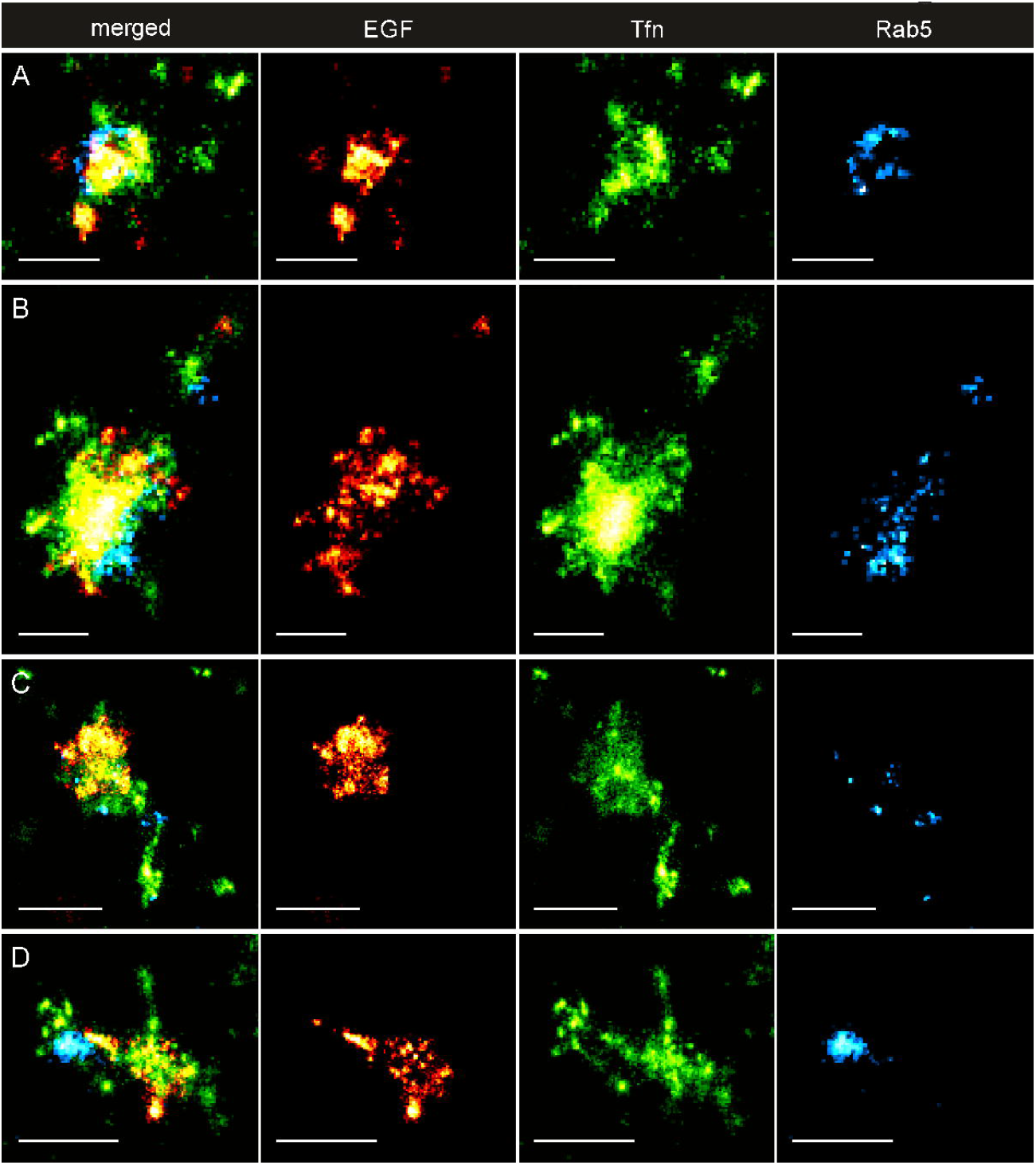
Triple-Colour Single-Molecule Localization Microscopy of endosomes in HeLa cells. Triple-Colour Single-Molecule Localization Microscopy of endosomes in HeLa cells reveals the compartmentalisation of early endosomes. A-D Representative examples of triple-positive endosomal structures displaying various types of compartmentalization of EGF (AF647, red), Transferrin (AF568, green) and Rab5c (Dronpa, cyan). EGF is predominantly present on a small part of the endosome, exhibiting a granular substructure. Transferrin signal spreads over the largest area of a presumed endosome and extends outwards in the form of elongated structures spanning several 100 nm. Its highest concentration are often exclusive of EGF hotspots. Rab5c is concentrated in nano- (A,C) and microdomains (B) on the endosomal membrane or in larger hotspots on the periphery (D). Scalebar: 250 nm (A,B), 500 nm (C,D).

Most strikingly, Dronpa-Rab5c was enriched in domains on the central part of the Tfn and EGF-positive structures. The Dronpa-Rab5c signals were found to line the apparent endosomal membrane in a circular geometry (Figure 2 A, Suppl. Figure S2A), separated from EGF and/or Tfn (Figure 2 B, Suppl. Figure S2B-G) or as small, distributed spots (Figure 2 C, Suppl. Figure S2B). Interestingly, Dronpa-Rab5c domains could also be found close to, or at the base of, elongated Tfn structures (Figure 2 C,D, Suppl. Figure S2C,E). To exclude misinterpretations due to possible photo-switching artefacts, we validated the SMLM imaging with multi-colour three-dimensional structured illumination microscopy (SIM), as distinct super-resolution method. We observed Dronpa-Rab5c and cargo geometries consistent with the SMLM images considering typical SIM resolution (Suppl. Figure S3).

We quantified the distributions of number and size of Rab5-domains in Tfn, EGF and Dronpa-Rab5c triple positive structures with a morphological particle analysis based on the SMLM images (see methods, Suppl. Figure S4). We determined on average 4.9 ± 3.1 (mean ± std) Dronpa-Rab5c domains with a diameter of 55.1 ± 37.8 nm (mean ± std) within the presumed endosomal area (Suppl. Figure S4A-C). In approximately 20 percent of these endosomes, we found only one, pronounced Dronpa-Rab5c domain with significantly enlarged diameter of 130.4 ± 46.3 nm (Suppl. Figure S4A). Interestingly, the total endosomal area covered by Dronpa- Rab5c follows a linear increase, dependent on the number of endosomal domains (Suppl. Figure S4D). Furthermore, while the coverage of endosomal area by Rab5c is virtually independent of the approximated overall size of endosomes (see methods, Suppl. Figure S4F), larger endosomes seem to exhibit more Rab5c domains (Suppl. Figure S4E). These results suggest that early endosomes have multiple Rab5-domains that may vary in number and size dependent on their size progression.

### A workflow for correlative multi-colour SMLM and electron tomography

We could resolve the two cargo molecules and Dronpa-Rab5c simultaneously with SMLM. However, since SMLM projects the three-dimensional endosomal structure to a two-dimensional image plain, it is not possible to unambiguously assign the fluorescent signals to the same or distinct endosomes (e.g. Figure 2 C, Suppl. Figure S2C-E). To address this limitation, we set to map the localization of molecules to the endosomal ultrastructure (e.g. limiting membrane of the central vesicle, ILVs, tubules). We established a correlative multi-colour SMLM and electron microscopy workflow (superCLEM) schematically represented in Figure 3. Hereby, the challenge was to take into account the structural specifics of endosomes as well as constraints of the sample preparation imposed by SMLM. Given the complex, vesiculo-tubular morphology of early endosomes, we used semi-thin (300-600 nm) sections to ensure that a major portion of the endosomal volume is contained within a section. In turn, to resolve the ultrastructure within semi-thin sections we had to apply electron tomography. Because heavy metal staining, dehydration and resin embedding are known to interfere with photo-switching of organic dyes and fluorescent proteins^45^, we chose to perform SMLM on thawed cryosections using the Tokuyasu method^46^. In contrast to standard resin embedding, this method enables sectioning of frozen hydrated samples and allows the staining with heavy metals after completion of SMLM imaging. Thereby, the Tokuyasu method yields better preservation of antigens, as well as fluorescence and photo-switching properties for SMLM^30^.

**Figure 3:**
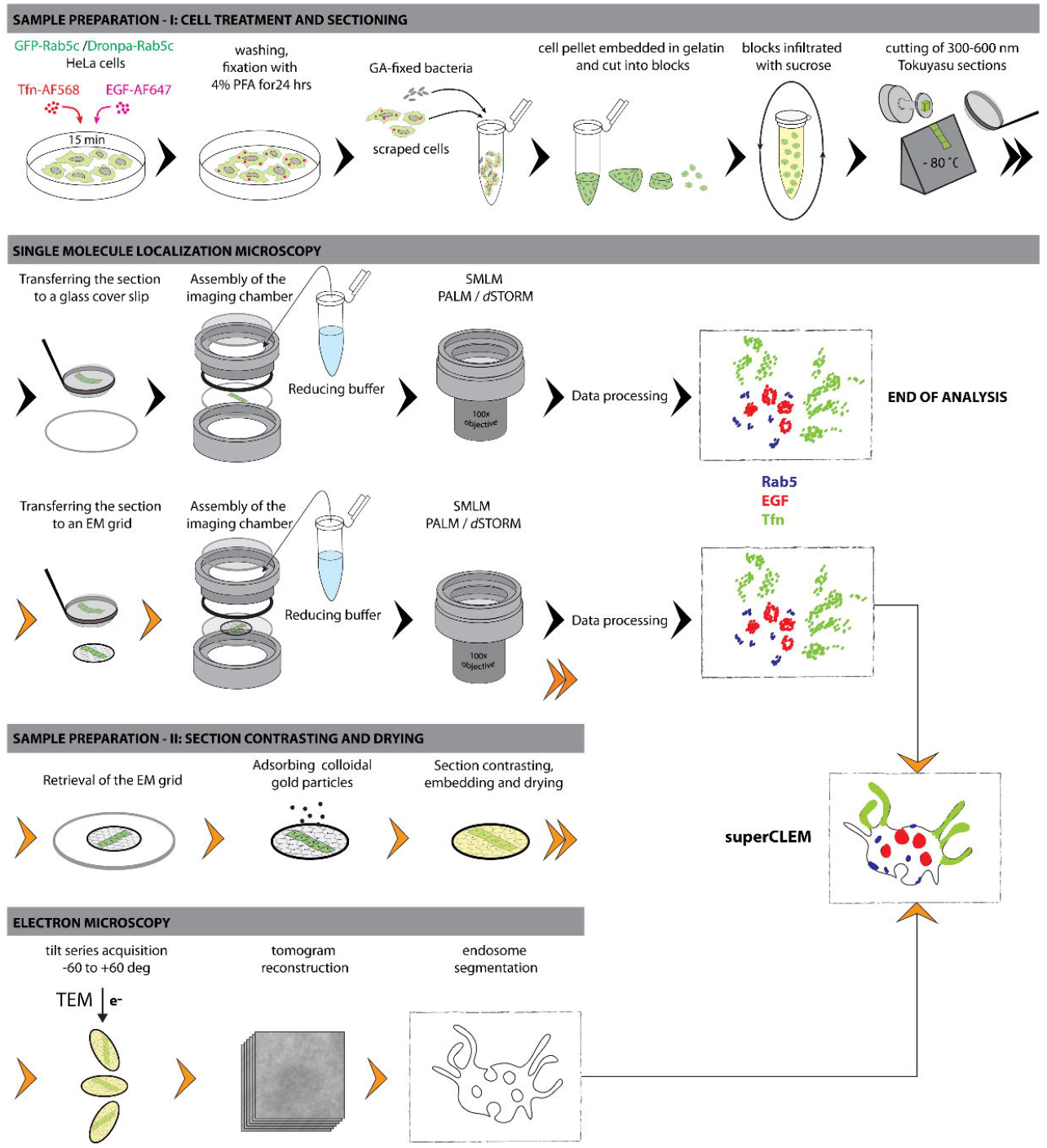
Schematic representation of the superCLEM workflow. Schematic representation of the workflow leading to an SMLM analysis with or without a subsequent electron microscopic analysis. Sample preparation and analysis are described in more details in the Results and in the Materials and Methods sections. Double arrowheads indicate that the workflow continues in a row below. Orange arrowheads indicate steps specific to the superCLEM approach.

For the final registration of the SMLM and EM data sets, we tested different types of fiducial markers visible in both imaging modes. Because beads that were adsorbed to the back side of the formvar film could obscure the region of interest (latex-beads) or cause strong out-of-focus background (tetraspecks) (data not shown), we ultimately chose glutaraldehyde (GA)-fixed bacteria as fiducials. These exhibit a strong broadband, photo-switching auto-fluorescence signal, and provide a stably immobilized marker within the actual imaging plain and restricted to the space between cells.

Our optimized superCLEM workflow is schematically represented in Figure 3. Briefly, after a 15-min pulsed uptake of cargo, cells were fixed, scraped, mixed with GA-fixed bacteria, pelleted and embedded in porcine gelatin. Subsequently, the pellet was cut into small blocks, which were infiltrated with sucrose to protect them from ice crystal formation when subsequently hardened by snap freezing in LN_2_ ^47^. Semi-thin sections were cut using a cryo-ultramicrotome and transferred either to glass cover-slips for SMLM imaging only, or to formvar-coated EM finder grids for the superCLEM approach. In the latter case, after SMLM imaging, the grids were washed, incubated with colloidal gold particles as EM-markers, contrasted with uranyl acetate, embedded in methyl cellulose and air-dried. Tilt-series of selected areas were acquired in a transmission electron microscope, followed by tomogram reconstruction and endosome segmentation. Finally, the SMLM image was aligned to the ultrastructural model to assign the super-resolved fluorescence information to the ultrastructural features.

### superCLEM - multicolour SMLM in the context of three-dimensional ultrastructure

Before applying the superCLEM protocol, we first ascertained that semi-thin sections of cells cut at random orientation retain the majority of endosomal structures by performing triple colour SMLM on sections deposited directly on glass coverslips (Figure 4, Suppl. Figure S5). We observed the same compartmentalization of the three markers in the sections as in whole cell SMLM, thus excluding significant distortions of observable structures. Moreover, because of the significantly reduced sample thickness, background and out of focus signals were virtually absent compared to whole cell imaging. Combined with the possibility to apply a full TIRF illumination, the detection probability and localization quality, especially with dimmer localization events were significantly improved, revealing fine-structures like thin Tfn extensions (Figure 4 A,C) and small EGF rings (Figure 4 A,D, Suppl. Figure S6). These results indicate that SMLM can be performed reliably on Tokuyasu sections.

**Figure 4:**
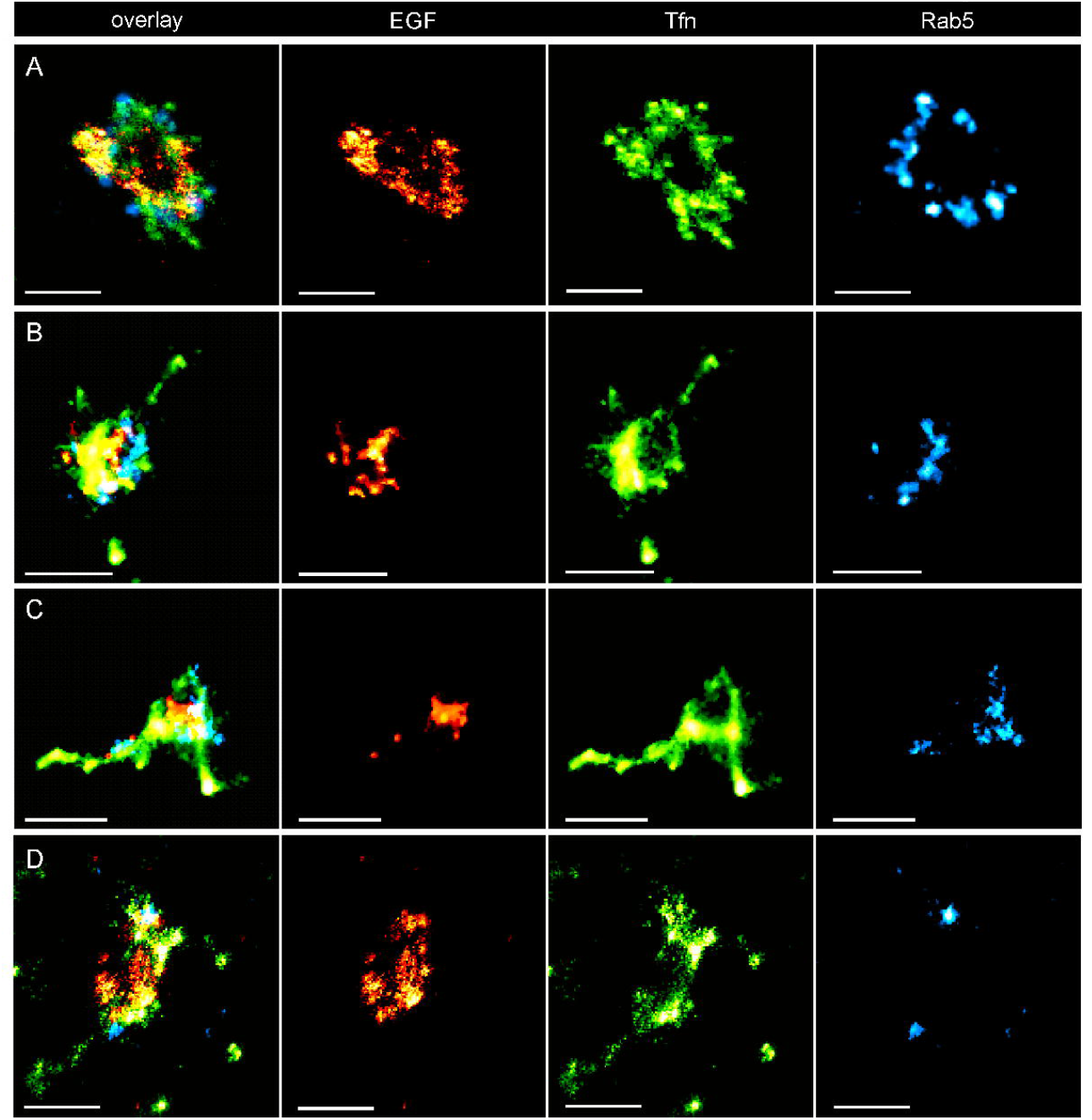
Triple-Colour SMLM of endosomes on Tokuyasu sections. Triple-Colour Single-Molecule Localization Microscopy of endosomes on Tokuyasu sections shows compartmentalization of early endosomes consistent with the analysis on HeLa cells. A-D Representative examples of endosomal structures displaying various types of compartmentalization of EGF (AF647, red), Transferrin (AF568, green) and Rab5c (Dronpa, cyan). Due to significantly reduced background and out of focus signal, the detection efficiency and quality is improved compared to whole cell SMLM. Thus, finer features like subtle Tfn tubules (A) and small EGF rings (A,D) can be resolved. Scalebar: 500 nm.

Next, we applied the superCLEM workflow to Tokuyasu sections deposited on EM finder grids. The SMLM data set in Figure 5 A shows the EGF and Tfn signals concentrated in distinct domains. After registration of the SMLM and the EM overview images (Suppl. Figure S7; exemplary), the tilt series of the selected region of interest (ROI) was acquired. The ultrastructural model was built by segmentation of membranes on virtual tomogram slices (Suppl. Figure S8, exemplary). The SMLM image of the ROI and the ultrastructural model were aligned (Suppl. Figure S9; exemplary) for a side-by-side comparison (Figure 5 A,B) to assign the fluorescent markers to distinct ultrastructural features of a single endosome. Whereas the EGF signal localized to the central vesicle, which is outlined by the limiting membrane, the Tfn signal mostly coincided with the tubular structures. A more in-depth analysis revealed that the spotty, sometimes ring-like SMLM EGF signal could be assigned to ILVs within the central vesicle. Two of these vesicles were not yet completely internalized, as their membranes were continuous with the limiting membrane of the central vesicle (Figure 5 C; magenta arrows). Most Tfn signal could be assigned to tubular structures, presumably recycling tubules, at one end of the central vesicle. Notably, the Tfn SMLM signal alone did not show a conspicuous tubular pattern due to the axial superposition of tubules, as shown by the EM analysis. Interestingly, some disperse Tfn signal was also apparent at the limiting membrane of the central vesicle, probably representing Tfn yet to be sorted into recycling tubules. One spot of Tfn signal appeared laterally close to tubules, but may correspond to a Clathrin-coated pit on the plasma membrane, axially distant from the endosome, as indicated by the side view of the ultrastructural model (Figure 5 B,C).

**Figure 5:**
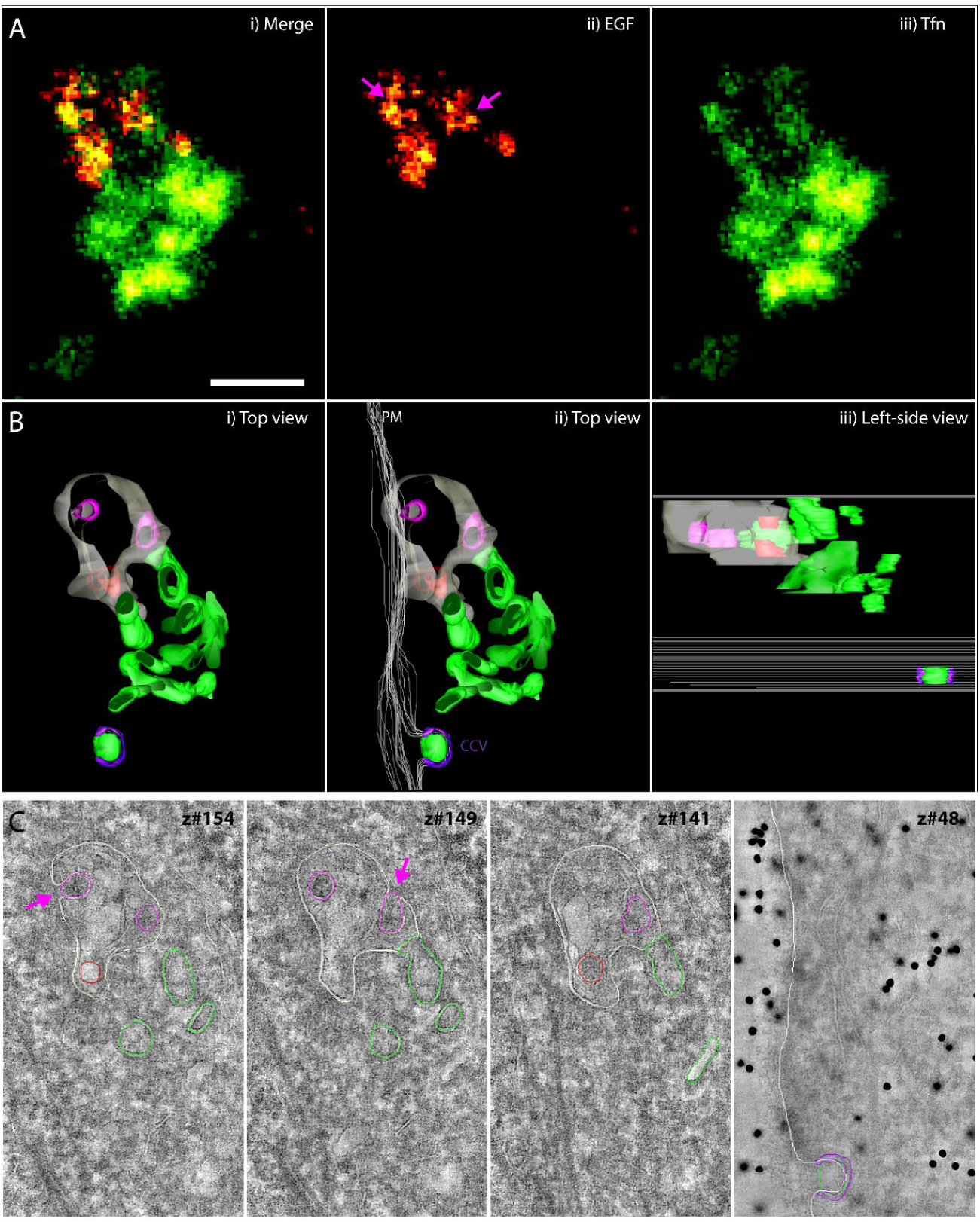
Visualization of EGF and Tfn cargo compartmentalization by superCLEM. (A) SMLM data for EGF-Alexa647 (red) and TFn-Alexa568 (green). (B) Ultrastructural model of the endosome based on a tomogram reconstructed from double-axis tilt series. (C) Virtual tomographic slices along four z-axis positions. Colour lines indicate segmented structures. Raw images are shown in Supple. Figure S8. Colours in B and C represent: (grey) limiting membrane, (red) ILV, (magenta) ILV continuous with the limiting membrane, (green) recycling tubules, (white) plasma membrane, (purple) clathrin coat. Pink arrows point to ILVs that are continuous with the limiting membranes and coloured magenta in the ultrastructural model. Scalebar: 250 nm.

Figure 6 provides examples that illustrate why the SMLM signals must be mapped to the ultrastructure of the organelle. When comparing the distribution of SMLM signals and ultrastructural models, EGF and Tfn are largely sorted to separate domains within the same endosome, e.g. in Figure 6 A. However, in the cases when their SMLM signals overlap, one could interpret that both are mingled and localized to the central vesicle (Figure 6 B,D). Yet, in such cases the EM analysis revealed the presence of tubular structures above or below the central vesicle, which can most likely be assigned to the Tfn signal (Figure 6 B,D). Furthermore, we identified many single Tfn positive structures, which when compared with the corresponding EM image did not appear to have a central vesicle (Figure 6 E). We assume these structures to represent neighbouring early or recycling endosomes. Particularly in the case where only a minor portion of an endosome was contained in a section the interpretation of the resulting sparse SMLM signal is challenging without ultrastructural context (Figure 6 C). Furthermore, although the EGF signal was generally localized to the central vesicle, it had an uneven distribution and, therefore, could not be considered a reliable marker to trace the outline of the central vesicle. In some reconstructed tomograms, we observed electron dense coats at the cytoplasmic side of the central vesicle (Suppl. Figure S8). Similar structures were previously described as sorting micro-domains, where EGF accumulates before internalization into ILVs^48,49^. We found many instances where, in addition to ILVs, the EGF SMLM signal could be assigned to sorting micro-domains, thus capturing different stages of EGF sorting (Figure 6 B, C, orange arrows).

**Figure 6:**
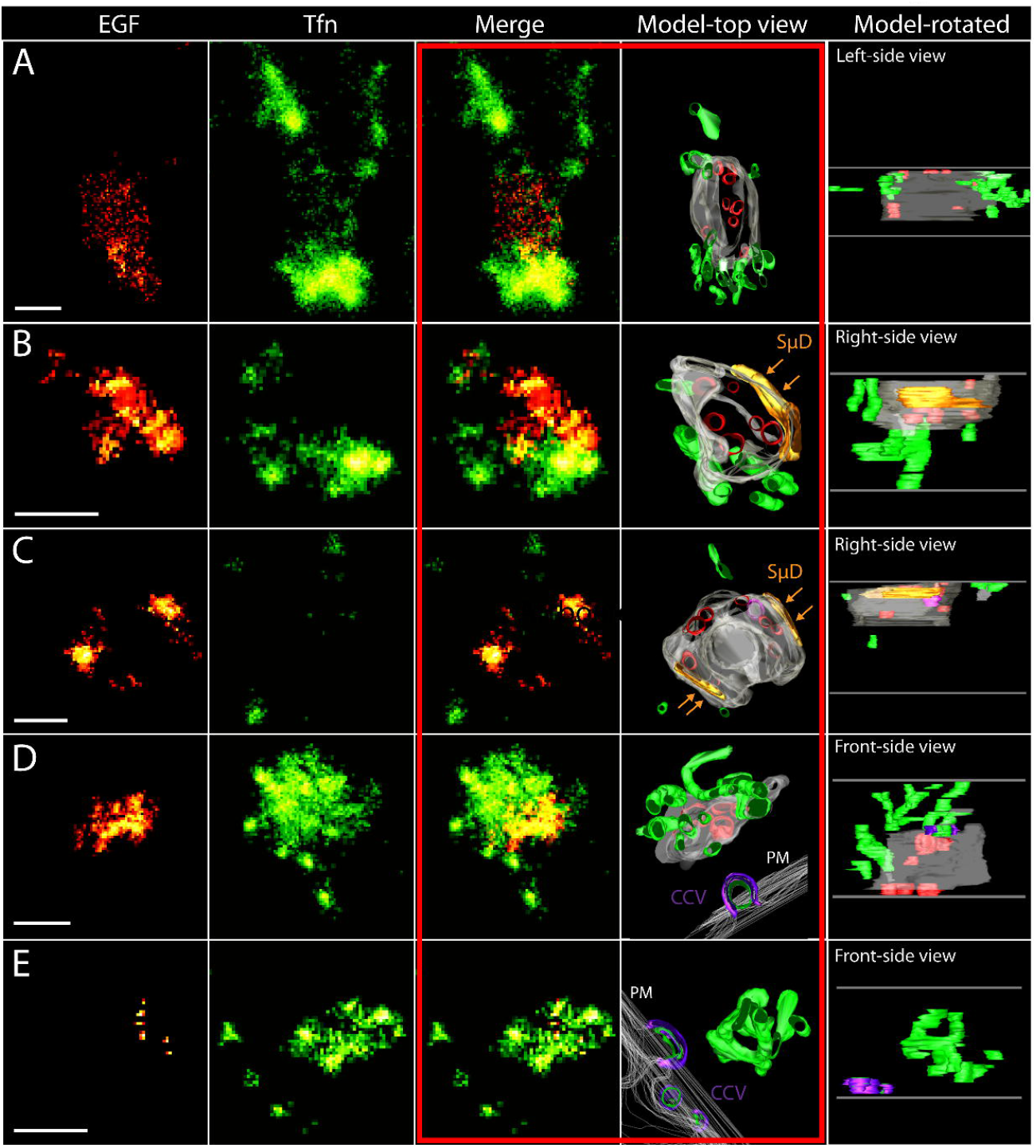
superCLEM reveals ultrastructural complexity behind SMLM projection images. Panels representing one particular endosome are arranged horizontally (A-E). SMLM data sets are presented in the first three columns: red colour is used to depict EGF-Alexa647 signal, green to depict Tfn-Alexa568 signal. Ultrastructural models are based on tomograms reconstructed from double-axis tilt series. Colours used in models represent: (grey) endosome limiting membrane, (green) tubular structures,(red) ILVs, (magenta) ILV continuous with the limiting membrane, (orange) sorting microdomains (SμD) on the limiting membrane. In panels D and E, in addition, plasma membrane (PM) was segmented and outlined in (white), clathrin-coated pits/vesicles are indicated with (purple) and the internalized cargo in (green) as it may coincide with the SMLM Tfn signal. Horizontal grey lines in the last column indicate top and bottom sides of a reconstructed tomogram. The third and fourth columns (framed with red to guide the eye) offer the optimal comparison between SMLM projection images and corresponding ultrastructural models. Scalebar: 250 nm.

The superCLEM workflow we developed can be applied to simultaneously assess the intra-endosomal distribution of multiple cargo molecules labelled with spectrally distinct organic dyes. We analysed the spatial distribution of EGF-Alexa647, Tfn-Alexa568 and LDL-Alexa488 internalized simultaneously by HeLa cells for 15 min. Whereas most Tfn SMLM signal could be assigned to complex tubular structures (Suppl. Figure S10), LDL and EGF signals could be mapped to the lumen of the central vesicle by EM analysis. However, in contrast to EGF that was concentrated in domains as described above, LDL exhibited a rather disperse pattern, which is consistent with the release from its receptor into the lumen^50^. We identified a number of ILVs within the globular part of the endosome, but only a fraction could be associated to the EGF SMLM signal (Suppl. Figure S6). Those ILVs most likely contain a distinct set of endocytosed membrane proteins that are destined for degradation. Furthermore, we performed SMLM on Tokuyasu sections with consistent localization patterns of all three cargo molecules compared to the superCLEM data set (Suppl. Figure S11)

Finally, we applied triple colour superCLEM to determine the localization of Rab5c (Dronpa- and GPF-Rab5c, see methods) in relation to the endocytic cargo. The results of representative superCLEM experiments are presented in Figures 7 and Suppl. Figure S12. Consistent with SMLM images acquired either on whole cells or on Tokuyasu sections deposited on cover slips, the Rab5c signal was localized in either multiple small (Figures 7 A, Suppl. Figure S12) or a singular larger domain (Figures 7 D, Suppl. Figure S12D). Expectedly Rab5c domains did not exhibit particular ultrastructural features. When the Rab5c SMLM signal was overlaid with the ultrastructural model, it largely coincided with the limiting membrane of the central vesicle (Figure 7 B,E; Suppl. Figure S12B,E). The analysis also showed that multiple Dronpa-Rab5c domains can be mapped to the same early endosome. In our superCLEM data sets we could also find instances, where Dronpa-Rab5c domains were located close to Tfn tubular bundles (Figure 7 B, Suppl. Figure S12B).

**Figure 7:**
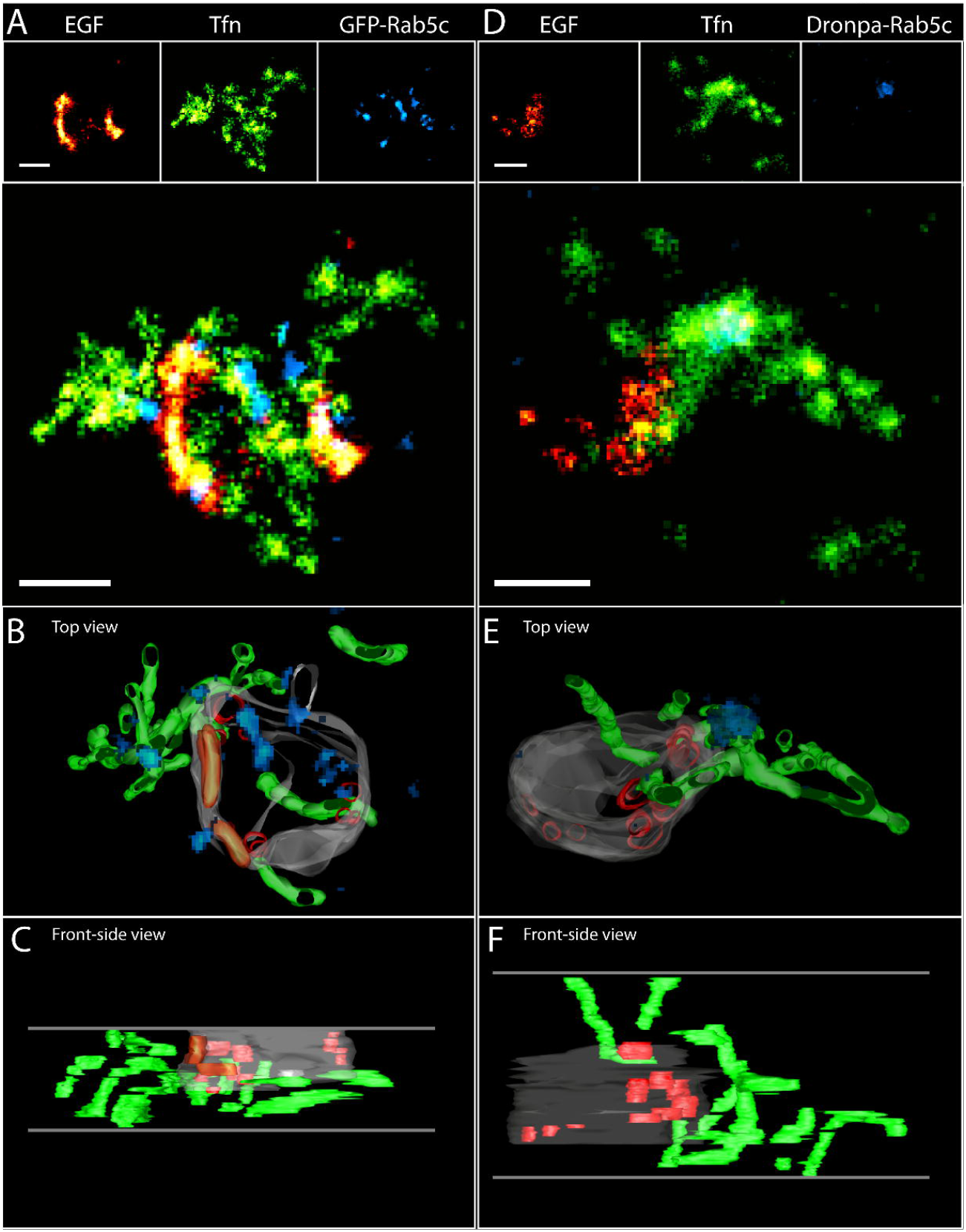
Mapping of Rab5c on endosomes visualized by triple-colour superCLEM. (A,D) SMLM data for EGF-Alexa647 (red), TFn-Alexa568 (green) and GFP/Dronpa-Rab5c (blue). (B,C,E,F) Ultrastructural models of endosomes based on tomograms reconstructed from double-axis tilt series. Colours used in models represent: (grey) limiting membrane, (red) ILV, (green) recycling tubules, (orange) sorting microdomains (SμD), (blue) Rab5c. Horizontal grey lines in the bottom images (C,F) indicate top and bottom sides of reconstructed tomograms. Scalebar: 250 nm.

## Discussion

In this study, we have addressed a so far unmet technical need to map several different proteins simultaneously to the ultrastructure of an early endosome with nanoscale resolution. For this, we developed a correlative triple-colour SMLM and electron tomography superCLEM workflow, combining *d*STORM and PALM on semi-thin sections, with three-dimensional ultrastructure imaging. To preserve the fluorescence properties of organic dyes and fluorescent protein after sectioning, we used a sample preparation protocol based on the Tokuyasu method. This way, it was possible to apply the heavy metal staining mandatory for EM after the completion of SMLM imaging.

We showed that semi-thin Tokuyasu sections mounted on glass cover slips are an optimal sample preparation for SMLM on organelles, due to the virtual absence of background and out of focus signal. Yet, the resolved fluorescence signal alone is not sufficient to make conclusive deductions about the overall endosomal structure or compartmentalization of cargo, even with SMLM resolution. This is mainly due to the difficulty to obtain reliably axially aligned three-dimensional SMLM ^51–53^ in a triple colour setup.

Our superCLEM workflow proved reliable with respect to previous notions of cargo sorting in the early endosomes. We could visualize different stages of EGF sorting within the globular part of early endosomes, such as micro-domains on the endosomal limiting membrane and formation of ILVs. In contrast, LDL was diffuse within the lumen of the central vesicle. Tfn was primarily located in recycling tubules with often complex, interwoven geometries, which are not resolvable by SMLM alone. These observations are in accordance with previously reported findings based on EM analyses^48,54–59^. However, neither EGF nor Tfn were exclusively confined to one particular compartment since we also found disperse Tfn signal on the limiting membrane of endosomes, and EGF signal neither associated with ILVs nor distinct micro-domains. This suggests that upon continuous internalization, we could detect cargo molecules transiently spread on the limiting membrane of a central vesicle prior to accumulation in endosomal domains. Thus, the SMLM protocol appears to be sufficiently sensitive to detect also sparse molecules on the membrane at various stages of cargo uptake and sorting in early endosomes.

Not every bit of the SMLM signal could be assigned to an ultrastructural counterpart. We conclude that during drying the fine ultrastructure at the very top and bottom of a section is less likely to be sufficiently preserved to allow segmentation. Furthermore, due to the missing wedge in electron tomography there is an inherent loss of structural information in certain regions of the tomogram. We compiled a detailed discussion of the limitations of our superCLEM approach and troubleshooting in the Supplementary Note.

In addition to resolving Tfn, EGF and LDL cargo, we found Dronpa-Rab5c localized to nano-domains within early endosomes. To our knowledge, this is the first report of a single-molecule based CLEM approach to precisely map different fluorescently-labelled proteins simultaneously to distinct domains within intracellular membrane compartments with a three-dimensional ultrastructural context. Based on the SMLM statistics performed on Tokuyasu sections deposited on glass coverslips, EGF and Tfn-positive endosomes have an average of five Dronpa-Rab5c domains with a diameter of ~55 nm, dependent on the overall size of the endosome. In all likelihood, these increase in number and size as they progress over time and ultimately undergo Rab5 to Rab7 conversion^60^. The oligomeric complexes of Rab5, its effectors and SNAREs previously detected by cross-linking and subcellular fractionation may underlie the formation of these domains^20^. Dronpa-Rab5c domains were also found associated with Tfn-positive recycling tubules. These results suggest that the biochemical cascades functionally linking Rab proteins in mammalian cells^61–63^ and yeast^64,65^ may also be spatially compartmentalized. Our superCLEM workflow will be instrumental to further dissect the molecular mechanisms underlying the structure, remodelling and function of endosomes.

Our method is generally applicable to the study of the sub-organelle organization at the nanoscale in other membrane compartments, like the Golgi apparatus, where the precise trafficking routes for cargo and resident enzymes are still debated^66^. The wide compatibility of our superCLEM workflow with photo-switchable proteins and organic dyes also enables other labelling approaches for SMLM. For example, expressing photo-switchable fluorescent proteins and self-labelling tags such as the SNAP- and/or Halo-tag^67^ utilizing the CRISPR/Cas9 system will allow the visualization of multiple proteins with superCLEM at endogenous expression levels. In addition, other cell permeable dyes that have been shown to be suitable for SMLM such as Mitotracker, ER-tracker, Lysotracker^68^; DAPI, Hoechst 33342^69^ could be used for superCLEM to label various organelles simultaneously. Further, the incorporation of fluorescence immunolabeling protocols on gently fixed whole cells before they are prepared for Tokuyasu sectioning, would significantly expand the potential of our approach, if sufficient ultrastructure preservation is maintained. Finally, adding a reliable approach for triple-colour three-dimensional SMLM to the current workflow will allow even better correlation between the two imaging modalities.

## Acknowledgements

We thank the MPI-CBG light microscopy facility for access and technical assistance especially Bert Nitzsche, Britta Schroth-Diez and Jan Peychl; the TransgeneOmics facility, especially Anne-Kristin Heininger, Ina Poser and Mihail Sarov for the design, generation, and maintenance of the BAC lines; the Electron microscopy facility, especially Tobias Fürstenhaupt for introduction to tomography; Ina Nuesslein, Julia Jarrells and Christina Eugster from the MPI-CBG FACS facility for assistance with the flow cytometry; Marc Bickle from the Technology Development Studio for assistance in the Yokogawa imaging. In addition, we thank Andreas Müller for valuable discussion and technical help with image registration. This work was financially supported by the German Research Foundation (DFG)(grant # 112927078, TRR 83 TP23), the European Research Council (ERC) (grant #695646 378) and the Max Planck Society (MPG).

## Material and Methods

### Cell culture

HeLa Kyoto cells were grown in DMEM supplemented with Penicillin (100 units/ml), Streptomycin (100 μg/ml) and Fetal Bovine Serum (10%). To generate a cell line expressing Dronpa-hRAB5C at near endogenous expression levels, the bacterial artificial chromosome (BAC) CTD-2005A24 was used to tag RAB5C with Dronpa or GFP (as described previously^70^) using BAC-TransGeneOmics.

A Dronpa::GGGGSGGGGS::Rab5c coding BAC was generated by Red/ET recombination from a Rab5c-containing BAC and a Dronpa-GS-linker PCR cassette. Homologous recombination yielded DNA containing tagged Dronpa-GGGGSGGGGShRAB5C, which was isolated and transfected into HeLa Kyoto cells using Effectene (Qiagen). We additionally transfected a Rab4a-BAC RP11-158E11 with a SNAP tag and generated it as mentioned above. Stably integrated transgenes were selected and maintained using media containing 400 μg/mL G418 (Invitrogen) and 90 μg/mL Nourseothricin sulfate. Three populations showing different levels of expression were sorted using a FACS Aria sorter.

### Western blotting

HeLa Kyoto were lysed using ice-cold lysis buffer (50 mM Tris, pH 7.5, 1% NP-40, 0.1% Na-deoxycholate, 150 mM NaCl, 1 mM EDTA + 1X Phosphatase Inhibitor Cocktail). For Western blotting equal amounts of HeLa cell extracts (10 μg) were subjected to SDS-PAGE and resolved, then wet-blotted on X Cell 2 Mini Cell apparatus (Invitrogen GmbH) onto nitrocellulose membranes, and subsequently incubated with primary antibodies. Rab5c was detected using a a rabbit primary antibody directed against Rab5c (Sigma HPA003426-100UL, 1/1000) was incubated with the membrane overnight at 4°C and, after washing with PBS-Tween buffer, incubated with an anti-rabbit HRP-conjugated secondary antibody (1/10,000, 30 min). Bands were detected with enhanced chemiluminescence Western blotting detection reagents (GE-Healthcare RPN 2106V1, 1:1, 1 min exposure time). Following the same procedure as for the Rab5c antibody, a mouse primary antibody directed against GAPDH (Sigma T6557-5ML, 1/1,000) was used on the same membrane to reveal GAPDH as a loading control. Densitometric quantification of Western blots was performed by using ImageJ software.

### Pulse-chase experiments

HeLa Kyoto cells and Dronpa-Rab5c or GFP-Rab5c cells from sorted populations were grown in 96-well microplates (GBO, #655866) for 24 h to reach 80-90% confluency. FBS-free media was added to the wells for one hour. CO_2_-independent medium was added to the wells for one more hour. Then at given time-points, the pulse solution (Tfn-Alexa568 [20 μg/ml], EGF-Alexa647 [1 μg/ml] in CO_2_-independent medium) was given to the cells. After one minute of pulse, the cells were washed in CO_2_-independent medium and then exposed to the chase solution (unlabelled Tfn [200 μg/ml] in CO_2_-independent medium). After the required time of chase, the cells were washed once in PBS and fixed in PFA 3% for 20 min and washed three times for 5 min in PBS. Then 50 μl of DAPI (1/5000) / Cell Mask Blue (CMB, 1/10000) in PBS were added to each well for 30 min. Finally, without removing the PBS/DAPI/CMB mixture, 90 μl of 0,04% sodium azide was added to each well. Imaging was carried out on a Cell Voyager CV7000 automated spinning disk microscope (Yokogawa) using a 40X/0.95NA objective. 8 fields of view (416 μm x 351 μm, 0.1625 μm/pixel) per well were acquired and conditions were tested in triplicates to yield 24 images per condition.

Image analysis was carried out using the lab custom built software Motion Tracking.

### Sample preparation for superresolution on glass

Coverslips were washed overnight with Hellmanex and then at least 3 times rinsed with water before they were stored in absolute ethanol. Before use, the ultra-clean coverslips (#1.5) (11 or 24 mm in diameter) were again intensively washed with water, dried and then coated with 50 μg/ml poly-L-lysine for 30 mins and then dried overnight and stored at 4°C. Before immediate use coverslips were washed 3x with PBS HeLa cells were seeded onto the coverslips at a desired density.

Depending on the application cells were exposed to a 15 min uptake of the ligand solution (Tfn-Alexa 568 [10 ng/ml, EGF-Alexa647 [100 ng/ml], Atto or Alexa 488-LDL [1:30 from home-made stock solution] in FBS-free DMEM). Cells were washed with 3x PBS and then fixed in 3% PFA for 15 min at 37°C and washed with PBS before SMLM imaging and for SIM imaging mounted with ProlongDiamond (Invitrogen).

### LDL purification and labelling

Low density lipoprotein (LDL) was purified from human plasma. The plasma was mixed with 0.22g of KBr/mL of plasma and subsequently centrifuged for 5 hours at 15°C. Then the mixture was overlaid with LDL-P (25 mM Na_3_PO_4_, 110 mM NaCl, 1 mM EDTA). The LDL fraction was then dialyzed twice against LDL-P + 5mM ascorbic acid. The protein yield was measured with the Pierce 660 assay and then the LDL solution was adjusted to pH=8.0 with phosphate buffer to then be coupled with the NHS-conjugated Atto or Alexa-488 dye for 1 hour at 25°C. To stop the reaction glycine was added for 10 min at RT and the labelled LDL was dialyzed then overnight against 5 L of PBS+50 mM ascorbic acid. After repurification through the same gradient as above the LDL-Atto/Alexa 488 was twice dialyzed ON as before.

### Structural illumination microscopy

For SIM, cells were imaged with a 60x, NA 1.42 oil objective on a Deltavision OMX v4 BLAZE (GE) equipped with 405, 488, 568, 642 nm lasers and four independent sCMOS cameras. Spherical aberration was minimized by choosing an immersion oil with a refractive index giving symmetrical point spread functions. Image stacks were acquired with 95 MHz, 0.125 μm z-steps and 15 images (three angles and five phases per angle) per z-section and a 3D structured illumination with stripe separation of 213 nm and 238 nm at 488 nm and 594 nm respectively. Image reconstruction was done using the Deltavision softWoRx 7.0.0 software with a Wiener filter of 0.0005-0.001 and wavelength specific OTFs specific for the mounting media.

### Single-Molecule Localization-Micoscopy

Image stack acquisition was performed with a *Nikon Eclipse Ti* microscope equipped with 640 nm, 561 nm, 488 nm and 405 nm laser lines. A 100X/1.49NA oil immersion objective together with a 1.5X post-magnification lens was used to achieve an optical pixel size of 104 nm. All measurements were performed with an active *perfect focus control.* In case of whole cell imaging a *HILO* illumination scheme was applied, whereas TIRF was utilized when imaging Tokuyasu sections. Typical acquisition parameters were 15-30 ms integration time per frame and 10000-30000 (647 nm), 20000-60000 (561 nm) and 5000-25000 (488 nm) acquisition frames total.

Although plain GFP is not widely used in SMLM it has been reported to exhibit intrinsic photo-switching upon 488 nm excitation and was applied to resolve nuclear patterns (Dickson et al., 1997; Gunkel et al., 2009). We therefore used it as a control reference for SMLM in case of superCLEM.

### Localization analysis

All raw image stacks were processed with rapidSTORM 3.2. For Tokuyasu sections the FWHM was restricted to typically 250 – 450 nm since no out of focus signal is expected, whereas for whole cell experiments the FWHM was typically restricted to 250 – 650 nm. Standard amplitude thresholds were set to 3000 (AF647), 1500 (AF568) and 450 (Dronpa, GFP) photons respectively. A framewise linear lateral drift correction was applied to localization sets and images were rendered in rapidSTORM to 10 nm lateral pixel size. In case of Rab5c, a post-localization median filter with 0-1 pixel radius was applied to reduce monomeric cytosolic signal.

The colour-channel overlay was performed either based on the bacterial fiducial markers or, in case of whole cell imaging, unambiguous endosomal structures (e.g. densely labelled, ring like features).

### Morphological analysis of endosomal domains

We selected the combined signals from cargo and Rab5 to approximate the total endosomal area without ultrastructural confirmation. We therefore performed a simple post-processing workflow in *ImageJ* based on the high-resolved images. A Gaussian-blurr with a radius of two pixel was applied to the multi-colour image before standard auto-thresholding (IsoData). The overall area was derived from the standard particle analysis including missing holes in the structure with a lower area threshold of 4 square-pixel. To quantify number and size of Rab5 domains, the Rab5 channel was analysed separately from the overall signal with a consistent workflow. Domain diameters were derived from the elliptical particle approximation. Histogramming, fitting and statistical analysis of domain data was performed with *OriginPro 2017*.

### Sample preparation for super-resolution CLEM

HeLa cells were exposed to 15 min uptake of the ligand solution (Tfn-Alexa 568 [10 ng/ml, EGF-Alexa647 [100 ng/ml], Atto or Alexa 488-LDL [1:30 from home-made stock solution] in FBS-free DMEM) at 37°C. Cells were washed with PBS 3x and then fixed with 4% PFA in 200 mM HEPES for 24 h at room temperature, protected from light. The fixative was removed, cells were washed with PBS and incubated with 0.1% glycine in PBS for 10 min to quench residual aldehyde fixative. Next, cells were scraped using 0.1% BSA in PBS and pelleted at 400 x g for 1 min. The cell pellet was resuspended in 12% porcine gelatin prepared in dH_2_O and mixed with GA-fixed bacteria resuspended in 12% porcine gelatin. Like HeLa cells, bacteria were extensively washed and incubated with 0.1% glycine after fixation. The mixture of HeLa cells and bacteria was pelleted at 4,500 x g, for 2 min. The tube was incubated on ice for 15 min to let gelatin solidify, then the pellet was removed and cut into small cubes of about 1-2 mm^3^. These were transferred to 2.3 M sucrose prepared in dH_2_O and incubated on a rotation wheel at room temperature at least overnight. For cryo-sectioning, a cube was mounted on an aluminium pin and snap frozen in LN_2_. A trim 45° and a cryo immunodiamond knives (Diatome, Switzerland) were used to trim a block and make 300-600 nm semi-thin sections at −80 °C, in an Ultracut EM UC6 ultramicrotome equipped with a cryo-chamber and a built-in EM CRION antistatic device (all Leica Microsystems). Using a mixture of 1.1 M sucrose and 1% methyl cellulose as a pick-up solution, sections were transferred onto cleaned 22-mm glass cover slips or onto H6 hexagonal pattern finder EM copper grids, coated with a formvar film on the pale side. Sections were stored at 4°C until imaged.

### Sample preparation for Single Molecule Localization Microscopy

For imaging a section deposited on a cover slip, the cover slip was clamped into an imaging chamber, which was then filled with water to dissolve a drop of methylcellulose/sucrose protecting the section from dehydration during storage. Next, the imaging chamber was filled with SMLM switching buffer and sealed with a second coverslip.

For imaging a section on an EM grid, a layer of methylcellulose/sucrose was dissolved by floating the grid on drops of cold dH_2_O. The grid was then soaked in a drop of 1% glycine in dH_2_O for 10 min to make the formvar film more hydrophilic and thereby prevent formation of a thin film of air, which would reflect light. Then an empty cover slip was clamped into an imaging chamber and the chamber was filled with SMLM buffer. The EM grid was transferred onto a cover slip in such a way that a side with a section was facing the cover slip. Only gentle force was used to push the grid against the cover slip, since during subsequent retrieval of an EM grid, we experienced difficulties with preservation of a formvar film when grids were tightly attached to a cover slip.

### Preparation of grids for electron tomography

Right after SMLM acquisition, the imaging chamber was disassembled. The coverslip mounted with the EM grid was transferred to dH_2_O. Carefully, the grid was removed from the cover slip and washed in several drops of dH_2_O. Next, the grid was incubated in a drop of 15-nm colloidal gold (Aurion, the Netherlands) for 2 min, to allow adsorption of gold fiducials onto both sides of the grid. After two more washes in dH_2_O, the back of the grid was carefully dried from the side using a small piece of filter paper and then the grid was floated on a drop of 0.2% uranyl acetate and 1.8% methyl cellulose for 5 min, on ice. The grid was looped out and air-dried.

### Electron tomography, reconstruction and registration

Double-axis tilt-series (−60 / +60 degrees) were acquired at 9,400x magnification on a Tecnai F30 transmission electron microscope operated at 300kV, using an UltraScan1000 CDD (2k x 2k) camera (Gatan, CA, USA) and the SerialEM software (Mastronarde, 2005). 3DMOD software (v. 4.9.8, University of Colorado, CO, USA) was used for tomogram reconstruction (using the weighted back-projection algorithm) and endosome segmentation. For segmentation the z-axis increment of 3 was applied to compensate for the collapse along a z-axis during a step of methyl cellulose embedding and drying of sections.

SMLM images were oriented with respect to low mag (2,300x-3,900x) montage EM overview images in the ICY software (v. 1.9.10.0, the Pasteur Institute, France) using bacteria embedded in the section as fiducials visible in both imaging modalities. The non-rigid transformation was applied for alignment. Tomograms were roughly aligned to the low mag overview EM image using distinct ultrastructural features (nuclear envelope, plasma membrane). In order to assign SMLM signals to an endosome model, the two datasets were compared when aligned side by side. To indicate endosomal compartmentalization, ultrastructural models were colour-coded. Green, representing Tfn, was used for tubules. Red, magenta, and orange, all representing EGF, indicated ILV, ILV continuous with the limiting membrane, and sorting microdomains, respectively. Selected examples of reconstructed tomograms and ultrastructural models are included as .mov files in the supplementary information.

