## Supplementary material for "Correlative SMLM and electron tomography reveals endosome nanoscale domains": Suppl.

**Supplementary Note**

Related to main text, providing additional information how to tackle superCLEM.

**Supplementary Information**

Figure S1: Validation of the GFP-Rab5c BAC HeLa cells. Related to Figure 1. Necessary validation of the additional cell line used.

Figure S2: Additional examples for triple-colour whole cell SMLM. Related to Figure 2. Relevant examples that illustrate consistency of results.
Figure S3: 3D-Structured illumination microscopy supports compartmentalization of Rab5c and cargo on endosomes. Related to Figure 2. Independent validation method to illustrate consistency of results.
Figure S4. Morphological analysis of Rab5c domains in triple positive endosomes. Related particularly to Figure 2 & 4.
Figure S5: Additional examples for triple-colour SMLM on Tokuyasu sections. Related to Figure 4. Relevant examples that illustrate consistency of results.
Figure S6: Distinct small EGF ring structures can be observed by SMLM on Tokuyasu sections. Related to Figure 4. Tokuyasu sections provide an optimal sample preparation for high-performance SMLM.
Figure S7: Registration procedure (overview). Related to Figure 3.
Figure S8: Examples of endosome segmentation. Related to Figure 3.
Figure S9: Registration procedure (endosome). Related to Figure 3.
Figure S10: Compartmentalization of EGF, Tfn and LDL on endosomes visualized by triple-colour superCLEM using semi-thin Tokuyasu sections. Relevant example that showcases the method.
Figure S11: Compartmentalization of EGF, Tfn and LDL on endosomes visualized by triple-colour SMLM. Relevant examples that illustrate consistency of method.
Figure S12: Mapping of Rab5c on endosomes visualized by triple-colour superCLEM (additional examples). Relevant examples that illustrate consistency of method.

Movie_Model_Fig-5:
Movie_Tomo_Fig-5:

Movie_Model_Fig-7i:
Movie_Tomo_Fig-7i:

Movie_Model_Fig-7ii:
Movie_Tomo_Fig-7ii:

Movie_Model_Fig-S10:
Movie_Tomo_Fig-S10:

Movie_Model_Fig-S12:
Movie_Tomo_Fig-S12:

**Supplementary Note**

### Our superCLEM workflow largely relies on a multi-step sample preparation, mainly to allow performing the heavy metal staining required for EM after the successful SMLM imaging. Along these steps we recognized a number of possible pitfalls leading to the destruction of the sample. Furthermore, due to the unique set of requirements demanded from the sample for superCLEM we think it to be important to elude on some finer details of the analysis.

Compared to the optimal case of Tokuyasu sections directly deposited on glass, sections applied to EM grids proved to be more challenging for SMLM imaging. This was because the formvar film supporting the sections cannot be completely flat, hence only a small region at a time was in focus. Additionally, because this setting often produced a slight gap between the section and the glass, we could not always apply a full TIRF illumination, leading to reduced overall SMLM quality. Furthermore, the copper EM-grid and the active contents of the photo-switching buffer, probably mainly the thiols, led to a progressing change in pH-value and thiol concentration during acquisitions, leading to suboptimal conditions for SMLM after the first measurement. We tried to circumvent this issue by using gold EM-grids but observed similar effects, probably due to progressing adhesion of thiols to the extended gold surface of the grids. Consequently, the SMLM quality, regarding photo-switching and single-molecule brightness, in superCLEM is slightly inferior compared to sections directly deposited on glass. For future endeavours the use of intrinsically blinking high-performance dyes could further improve the method.

Drying the back-side of a grid with the filter paper after SMLM imaging is probably the most delicate step of the whole protocol with great impact on the ultrastructural preservation. If the front side of a section is accidentally exposed to drying (in the absence of methylcellulose), its ultrastructure will be irreversibly destroyed. Additionally, the formvar film can get torn during this procedure. Especially during the early stages of the protocol development we lost a significant portion of samples around this step. We assure potential users that with some patience and practice this step can be performed with sufficient success rates, although a certain loss ratio will remain.

Due to intrinsic restrictions of the individual methods may cause that signals cannot fully or not certainly be assigned to an underlying ultrastructure. These restrictions include the finite angular range (‘missing wedge’) in electron tomography, shrinkage of sections after drying and the correlation of 2D fluorescence data with 3D electron tomography.

We experience the missing wedge as a problem especially for tubular structures, as it hard to observe their connections with the central vesicles due to the limited information that can be acquired by tomography.

On the other side, we could also observe tubular structures associated with the central vesicle of an endosome that did not have an obvious counterpart in the SMLM image (e.g. Supplementary Figure 6). Given the density of SMLM Tfn signal in tubules, these most likely represent other recycling tubules that contain different cargos or tubules that are destined for retrograde transport.

We observed a significant axial shrinkage of sections after drying, estimating a remaining 1/3 of their initial thickness. This allowed us to analyse sections that were originally thicker than the commonly used 300 nm, though it is likely that some distortions to the ultrastructure took place. Tokuyasu sections are so far not frequently used for electron tomography and the structural collapse along a z-axis is probably the main reasons. On the positive side, the membranes are well preserved and appear clearly white due to the negative staining. As pointed out in the main manuscript, the delicate structures of tubules originating from the globular part of endosomes are consistent with resin embedding, where no drying and the connected axial shrinkage can take place. So we are confident that shrinkage will lead to ultrastructural compression rather than major distortion, but users have to keep in mind that this may not be the case for alternative target structures.

Finally, the correlation of two-dimensional SMLM data to the three-dimensional EM tomograms is only possible under certain conditions. The usage of densely seeded fiducial markers in the imaging plane allows the mapping of SMLM data to ultrastructure with nanometer precision laterally, but not in axially, where signals can be assigned to certain likely structures. Concordantly, based on *a priori* knowledge about the proclivity of Tfn to reside in recycling tubules and EGF in ILVs we experience that after endosome segmentation particular cargo could be assigned to particular compartment with certain probability. Adapting the workflow to three-dimensional SMLM will allow a complete correlation between imaging modi, but potential users should be aware of chromatic shifts and other artefacts that complicate multi-colour three-dimensional SMLM.

**Supplementary Information**

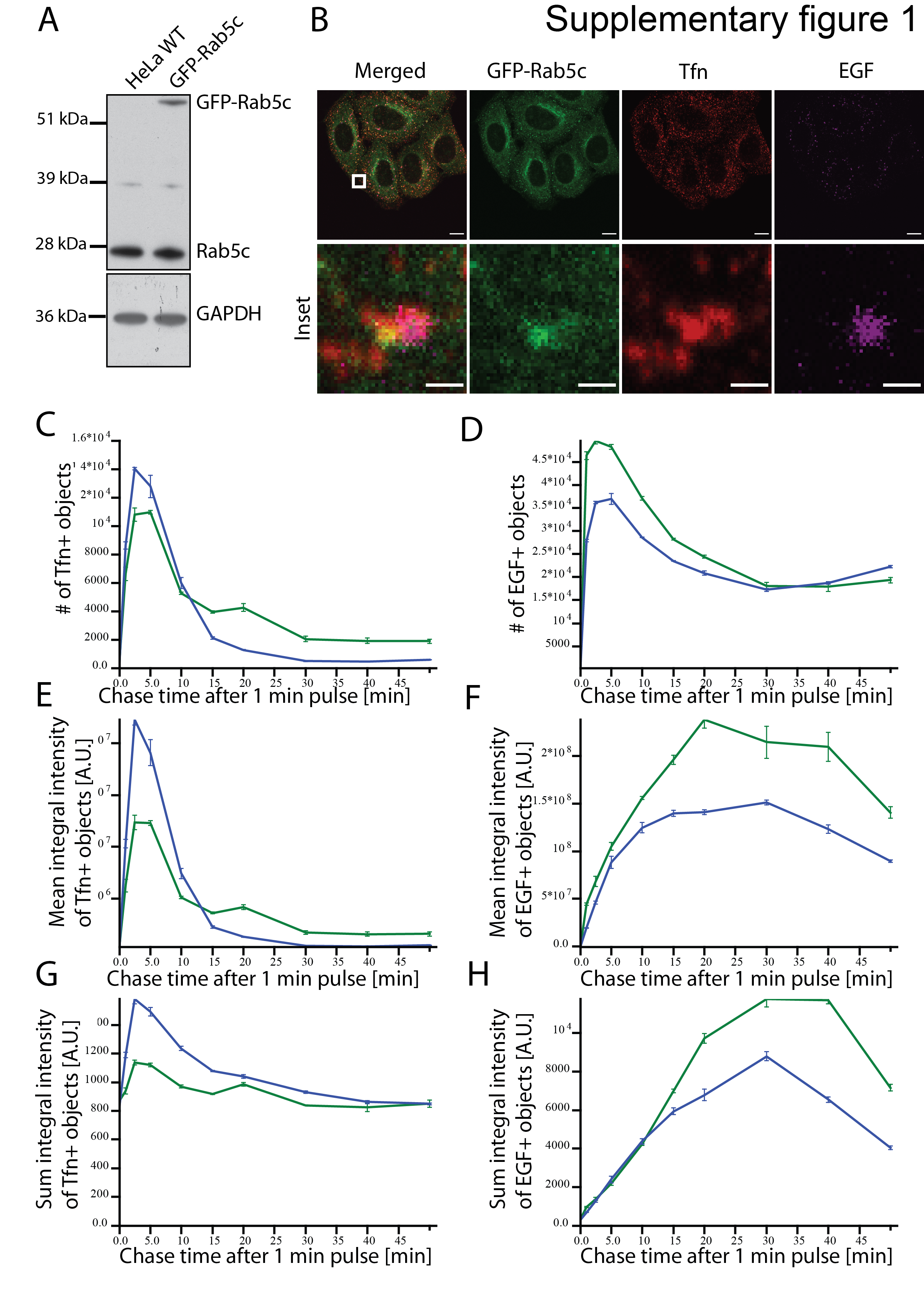

**Figure S1: Validation of the GFP-Rab5c BAC HeLa cells.**

(A) Immunoblot of GFP-Rab5c cells with high GFP-Rab5c expression with an anti-Rab5c antibody. The GFP-tagged protein has an expected size of ~50 kDa. The percentage of each population was 30% relative to the total Rab5c amount. The GAPDH band was used as loading control.

(B) Immunofluorescence of GFP-Rab5c cells. Cells were fed with Tfn and EGF continuously for 15 min before fixation. Displayed is a representative image of the cells (upper panel) with an Inlay (lower panel) that is the enlargement of the white rectangle. Confocal images were analysed using the Fiji software. Scale bar: 10 µm, Inlay: 1 µm

(C-H) Kinetics of Tfn (C, E and G) and EGF (D, F and H) trafficking by the GFP-Rab5c BAC cell line (green) and control HeLa (blue) (Mean ± SEM)

(C-D) Number of Tfn- (C) and EGF-positive (D) endosomes per masked area.

(E-F) Mean integral intensity of Tfn- (E) and EGF-positive (F) endosomes per masked area.

(G-H) Sum integral (overall) intensity of Tfn- (G) and EGF-positive endosomes (H) per masked area.

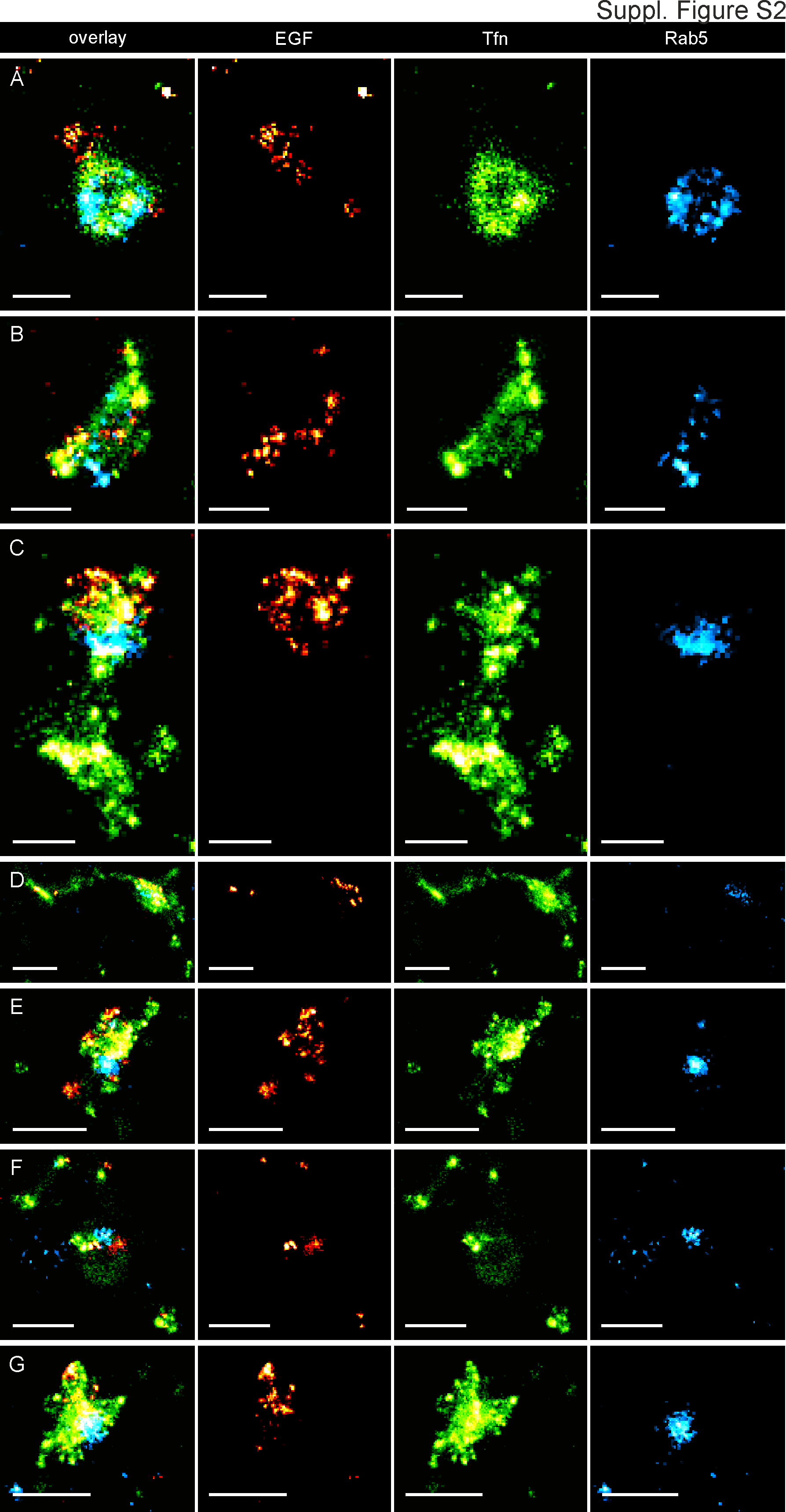

**Figure S2: Additional examples for triple-colour whole cell SMLM**

Triple-Colour Single-Molecule Localization Microscopy of endosomes in HeLa cells reveals the compartmentalisation of early endosomes. A-D Representative examples of endosomal structures displaying various types of compartmentalization of EGF (AF647, red), Transferrin (AF568, green) and Rab5c (Dronpa, cyan). Scalebar: 250 nm (A-C), 500 nm (D-G).

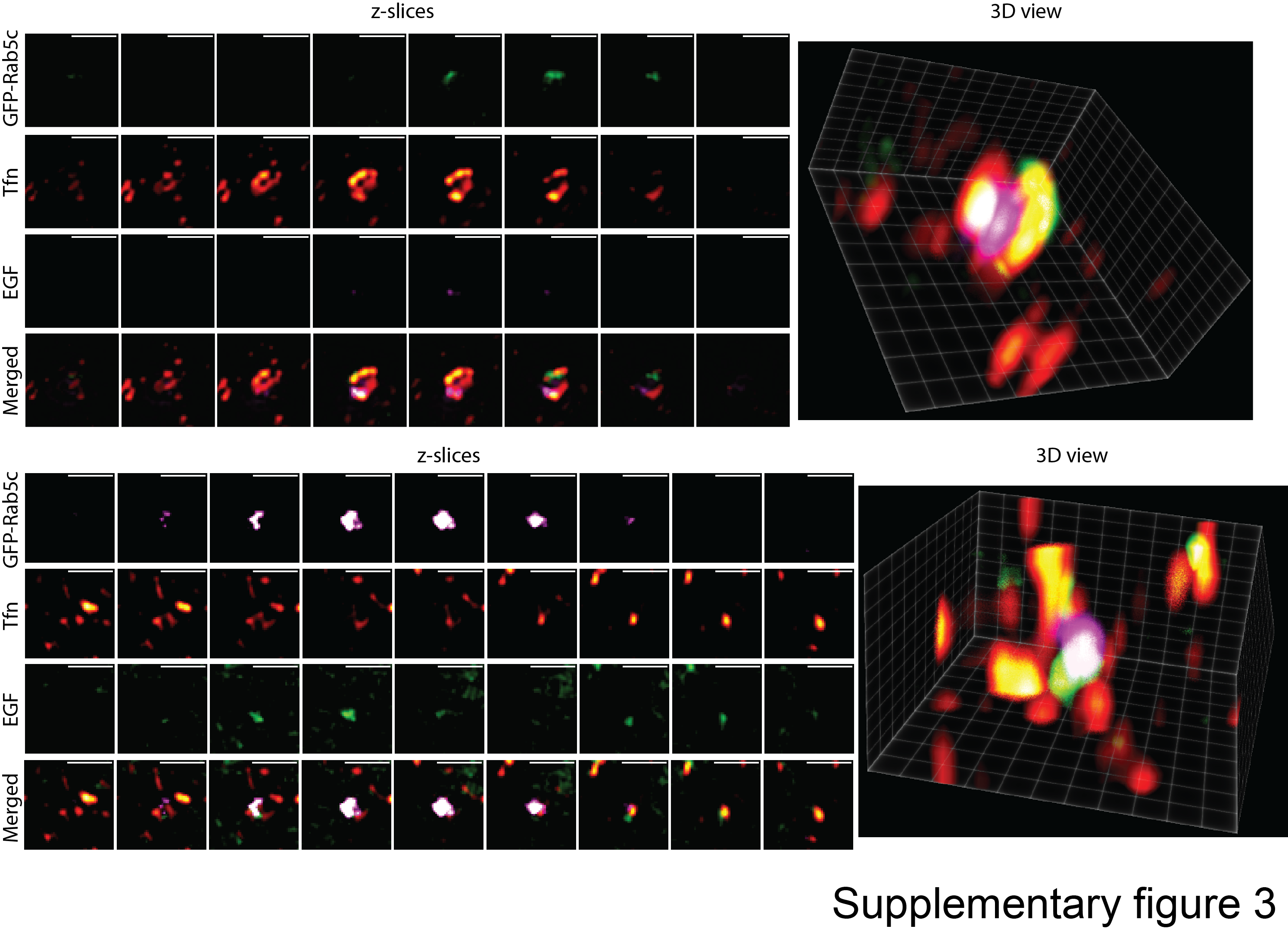

**Figure S3: 3D-Structural illumination microscopy supports compartmentalization of Rab5c and cargo on endosomes.**

3D SIM characterization of Rab5c early endosomes positive for Tfn and EGF. GFP-Rab5c-labelled were fed with Alexa-568 Tfn and Alexa-647 EGF for 15 mins, fixed and then analysed with SIM. Individual endosomes marked by a white rectangle were enlarged and individual channels are shown in the direction of the plasma membrane facing the coverslip upwards. Cytosolic background staining from GFP-Rab5c are a consequence of inactive Rab5c residing in the dense perinuclear area. Scalebar: 10 μm and 1 μm in the enlargements.

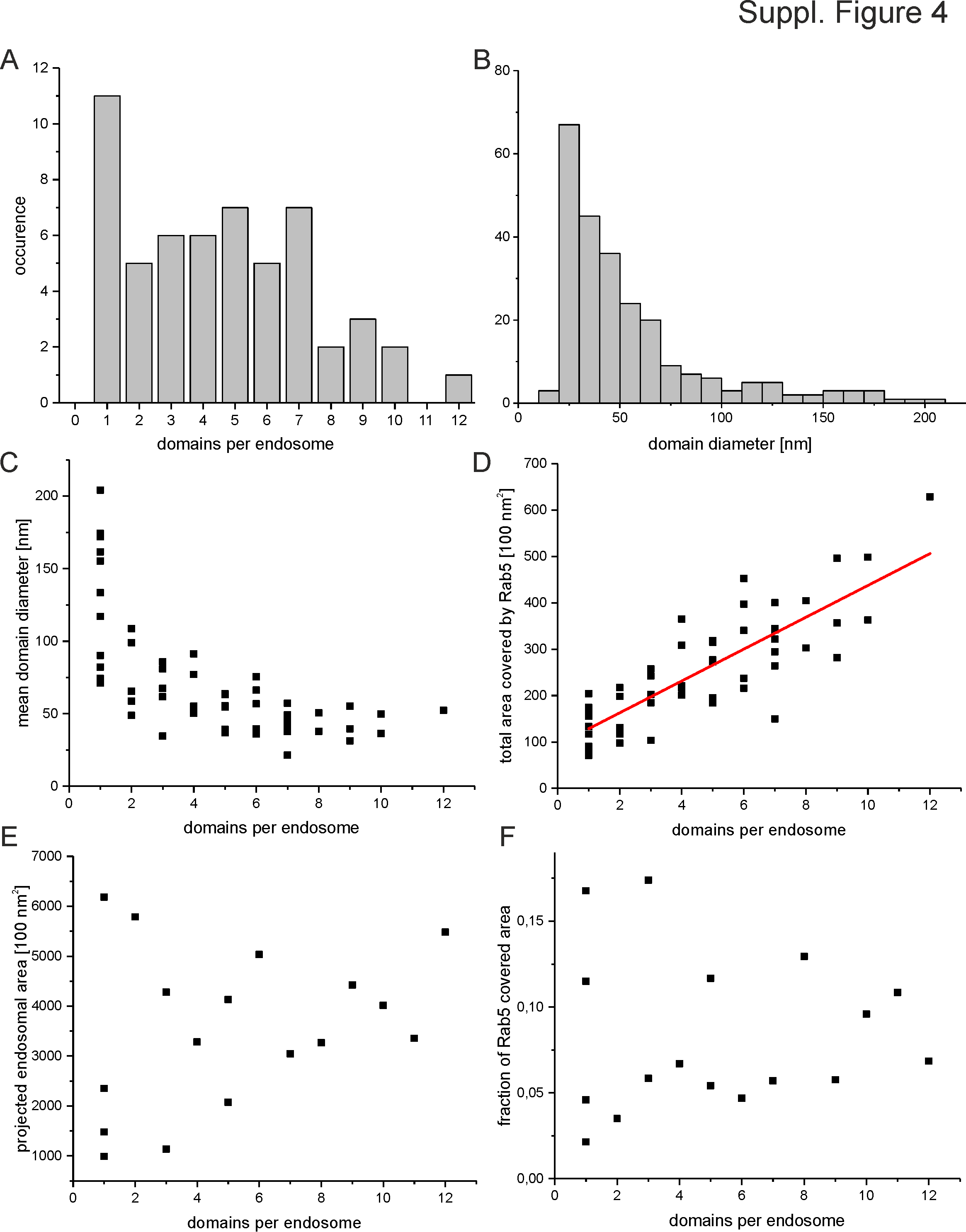

**Figure S4. Morphological analysis of Rab5c domains in triple positive endosomes.**

(A) Number of Rab5c domains per triple-positive endosome (4.9+-3.1, mean+-std).
(B) Distribution of Rab5c domain diameter. (55.1+-37.8nm, mean+-std)
(C) Mean Rab5c domain diameter dependent on the number of Rab5c domains within the endosome
(D) Total endosomal area covered by Rab5c domains dependent on the number of Rab5c domains within the endosome

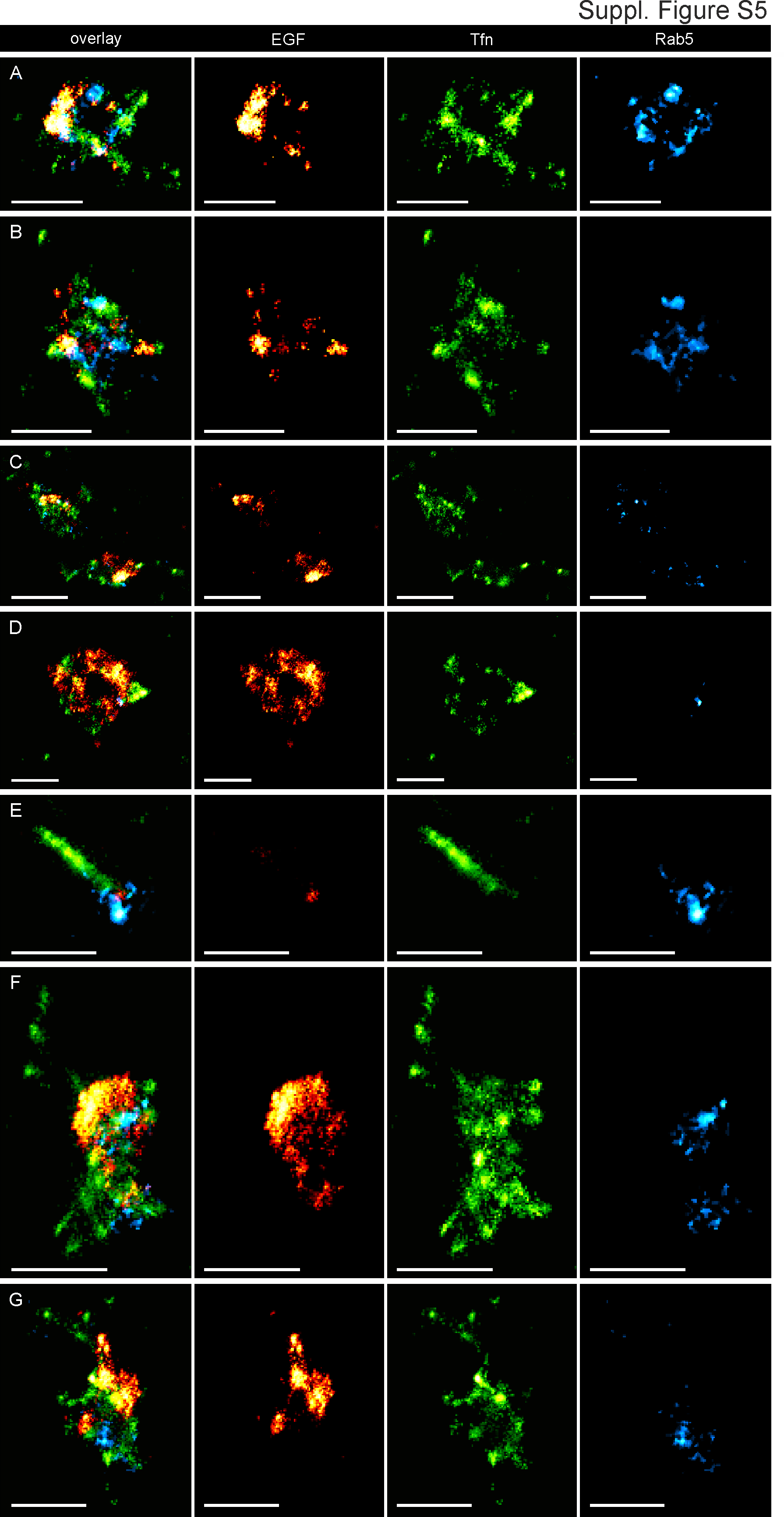

**Figure S5: Additional examples for triple-colour SMLM on Tokuyasu sections**

Triple-Colour SMLM of endosomes on Tokuyasu sections shows compartmentalization of early endosomes consistent with the analysis on whole cells. A-D Representative examples of endosomal structures displaying various types of compartmentalization of EGF (AF647, red), Transferrin (AF568, green) and Rab5c (Dronpa, cyan). Scalebar: 500 nm. **Figure S6: Distinct small EGF ring structures can be observed by SMLM on Tokuyasu sections.** Original areas from zoom images are indicated as white boxes, while line-profile traces are indicated as white arrows. Black squares in line-profiles indicate data points, red, solid lines bimodal fits to the data. The peak-to-peak distances are indicated as coloured values in every line-profile. Scalebar: 250nm.

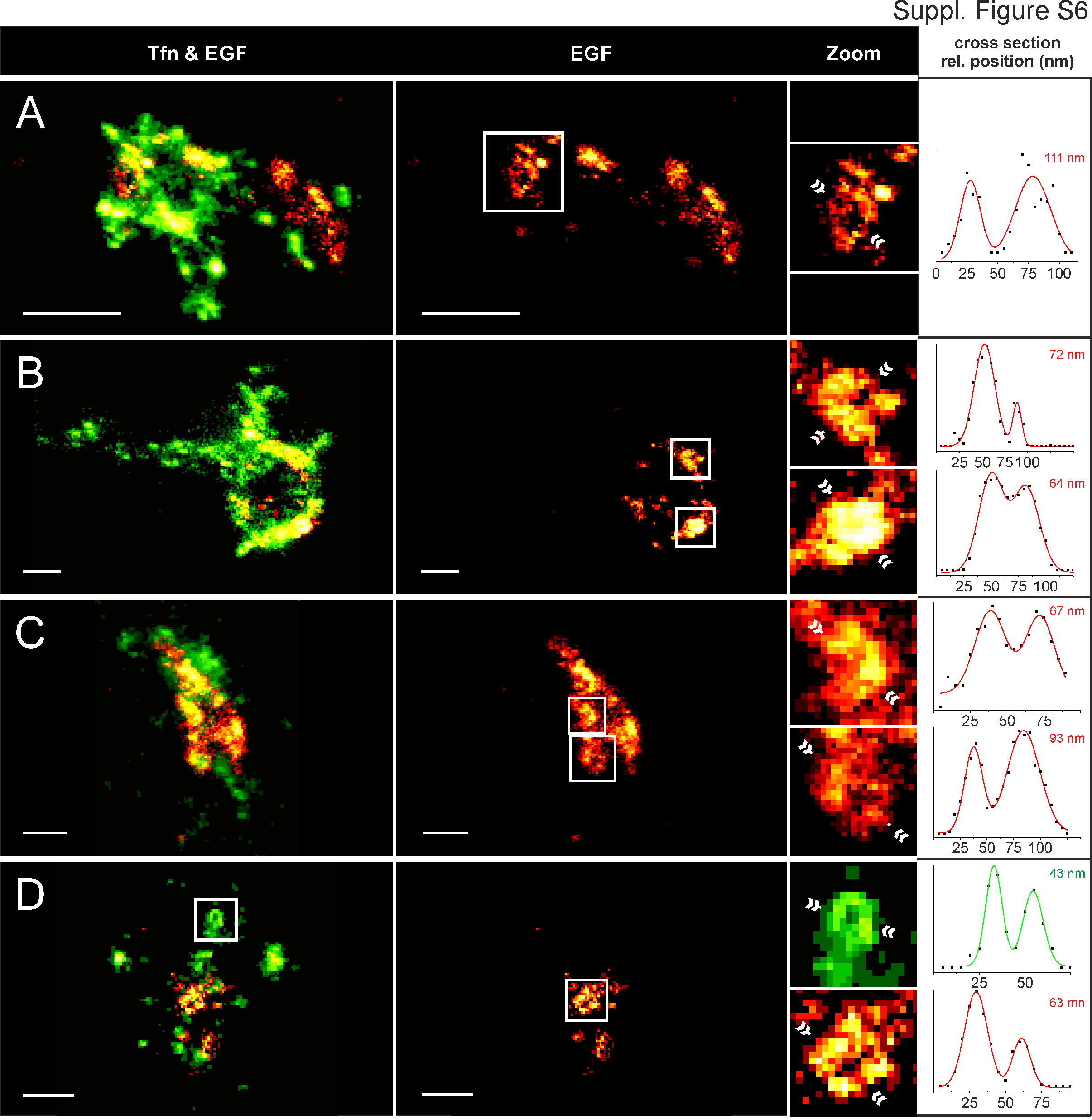

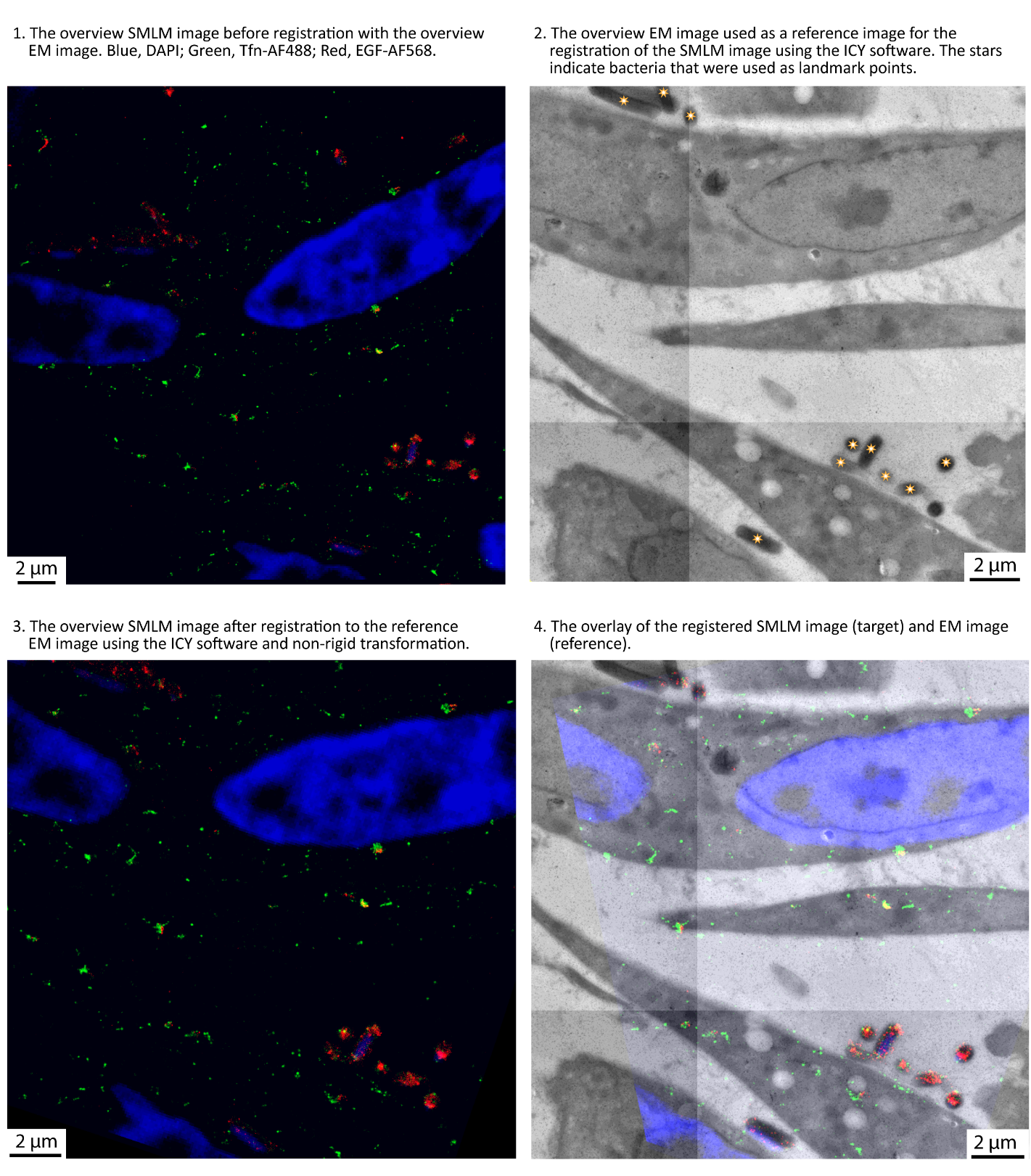

**Figure S7: Registration procedure (overview).**

Illustration of the registration procedure of the SMLM and the EM overview images using fluorescent bacteria (indicated with stars on the upper right image) as fiducials and the ICY software.

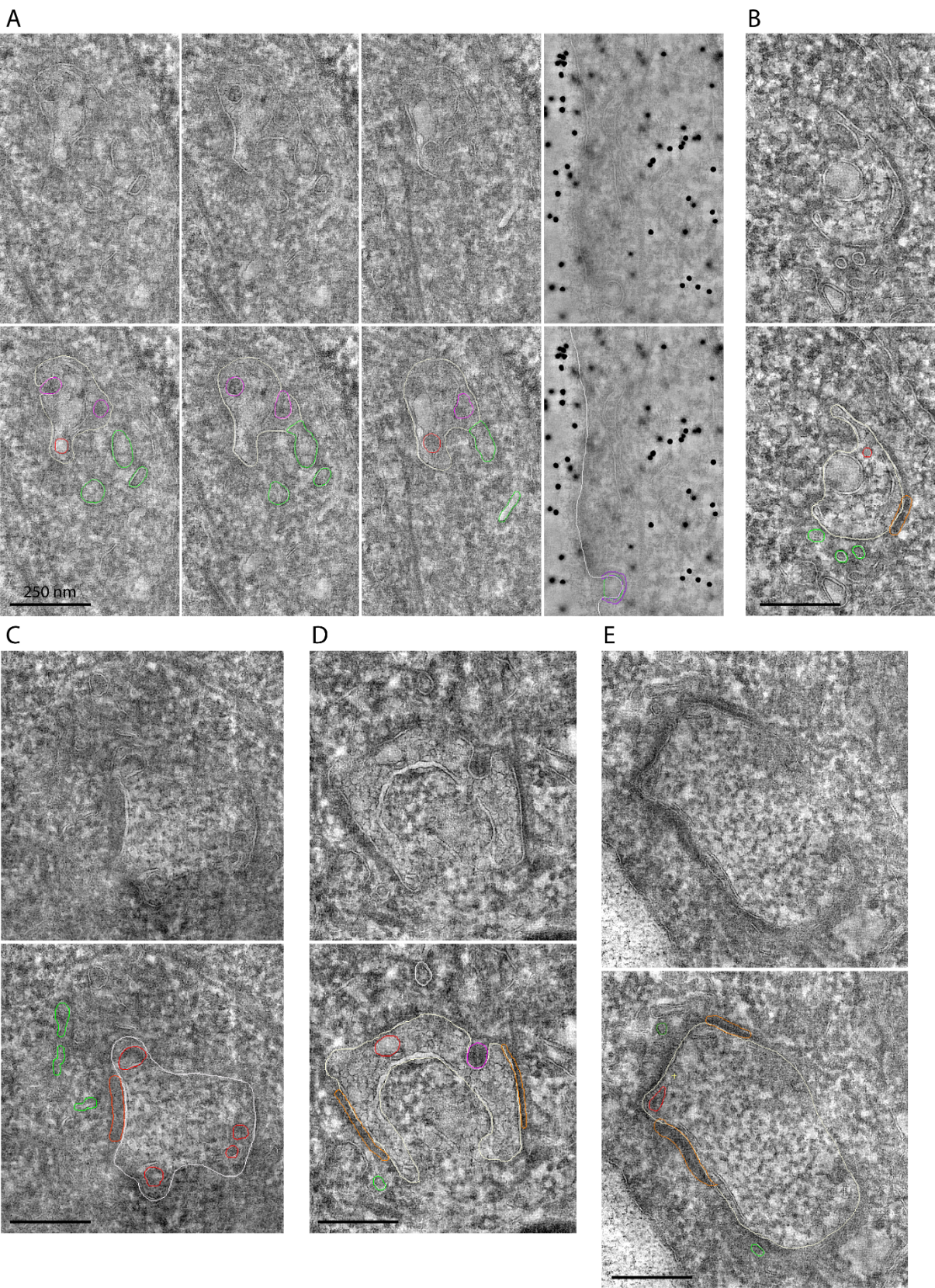

**Figure S8: Examples of endosome segmentation** in five different tomograms (A-E). Colour lines indicate structures that were segmented: (grey) endosome limiting membrane, (green) tubular structures, (red) intraluminal vesicles - ILVs, (magenta) ILV continuous with the limiting membrane, (orange) sorting microdomain on the limiting membrane, (purple) clathrin-coated vesicle.

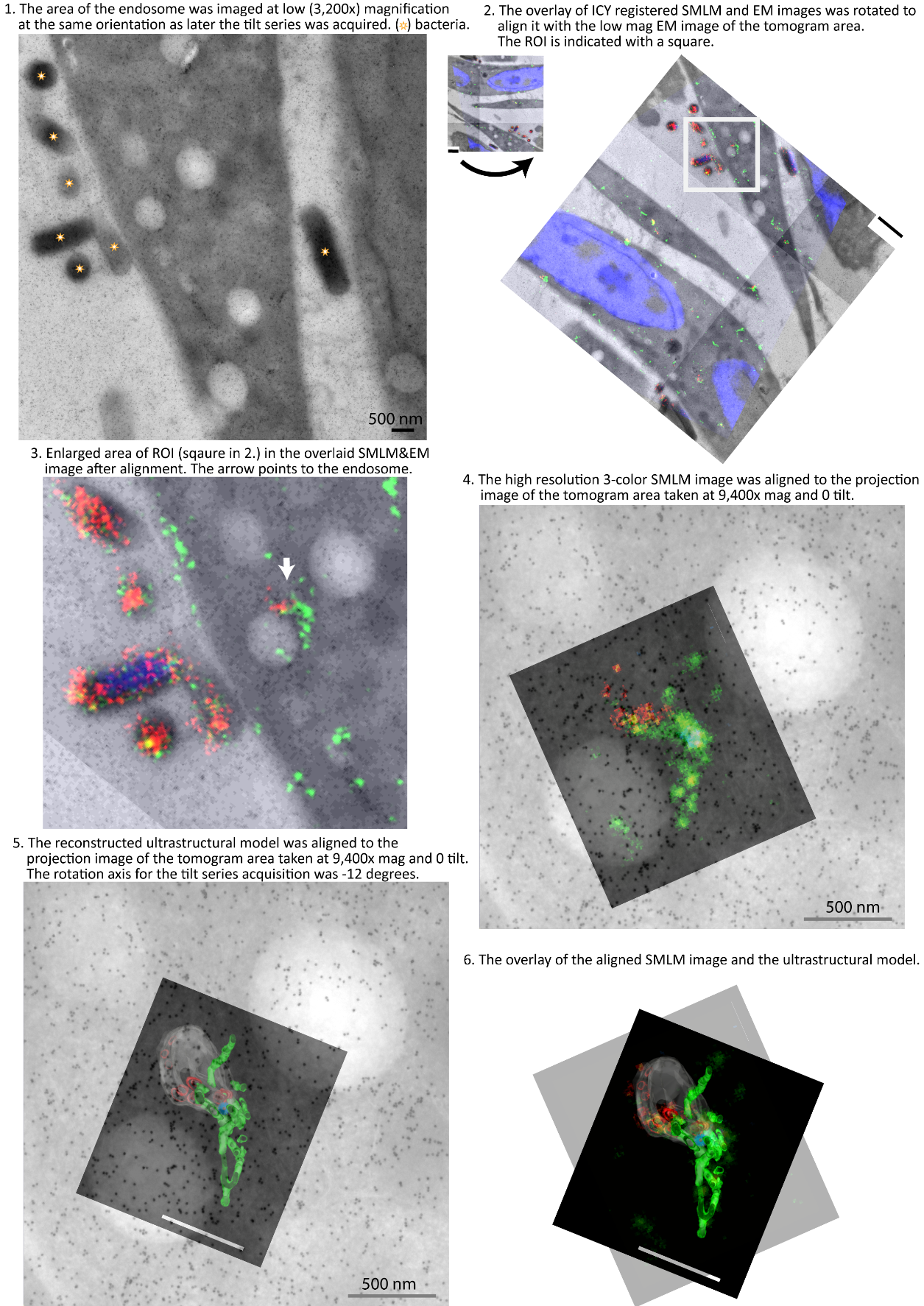

**Figure S9: Registration procedure (endosome).**

Illustration of the alignment procedure of the SMLM image and the ultrastructural model.

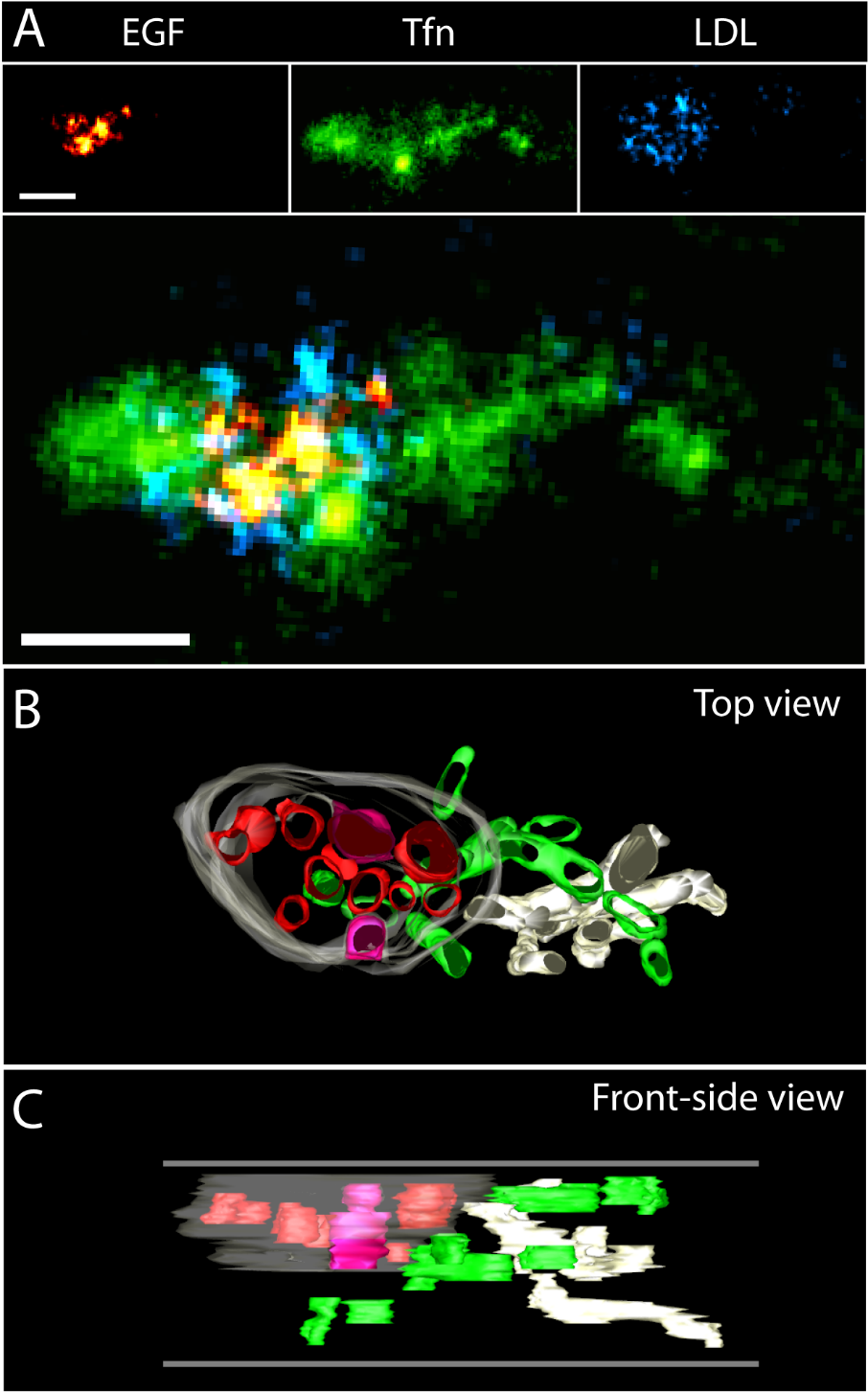

**Figure S10: Compartmentalization of EGF, Tfn and LDL on endosomes visualized by triple-colour CLEM using semi-thin Tokuyasu sections.**

(A) SMLM data for EGF-Alexa647 (red), TFn-Alexa568 (green) and LDL-Alexa488 (blue). (B,C) Ultrastructural models of the endosome based on a tomogram reconstructed from double-axis tilt series. Colours used in models represent: (grey) limiting membrane, (red) ILV, (magenta) ILV continuous with the limiting membrane, (green) recycling tubules. LDL is localized to the lumen of the central vesicle. Horizontal grey lines in (C) indicate top and bottom sides of a reconstructed tomogram. Scalebar: 250 nm.

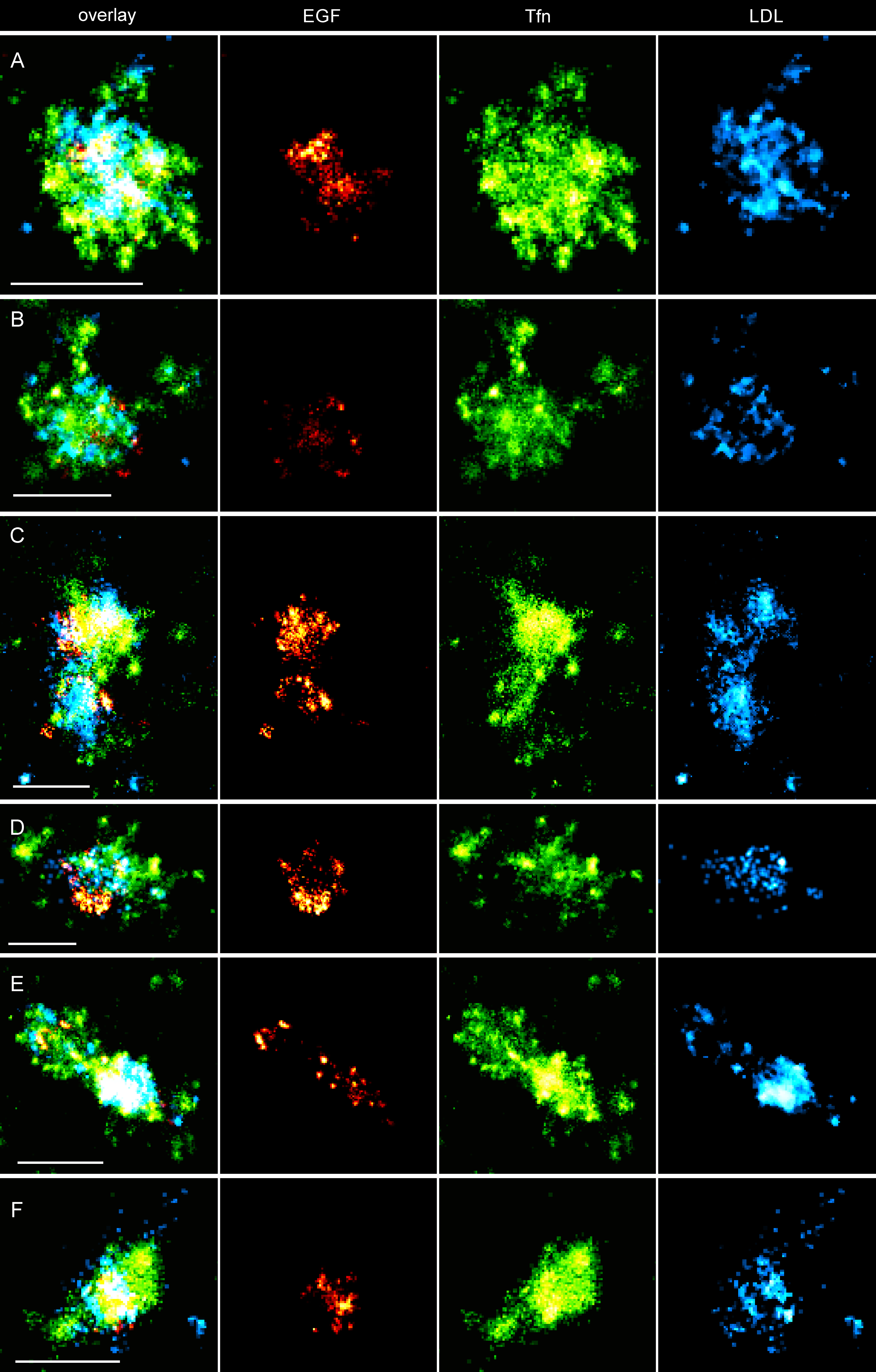

**Figure S11: Compartmentalization of EGF, Tfn and LDL on endosomes visualized by triple-colour SMLM.** EGF-Alexa647 (red), TFn-Alexa568 (green) and LDL-Alexa488 (cyan). Scalebars: 500 nm.

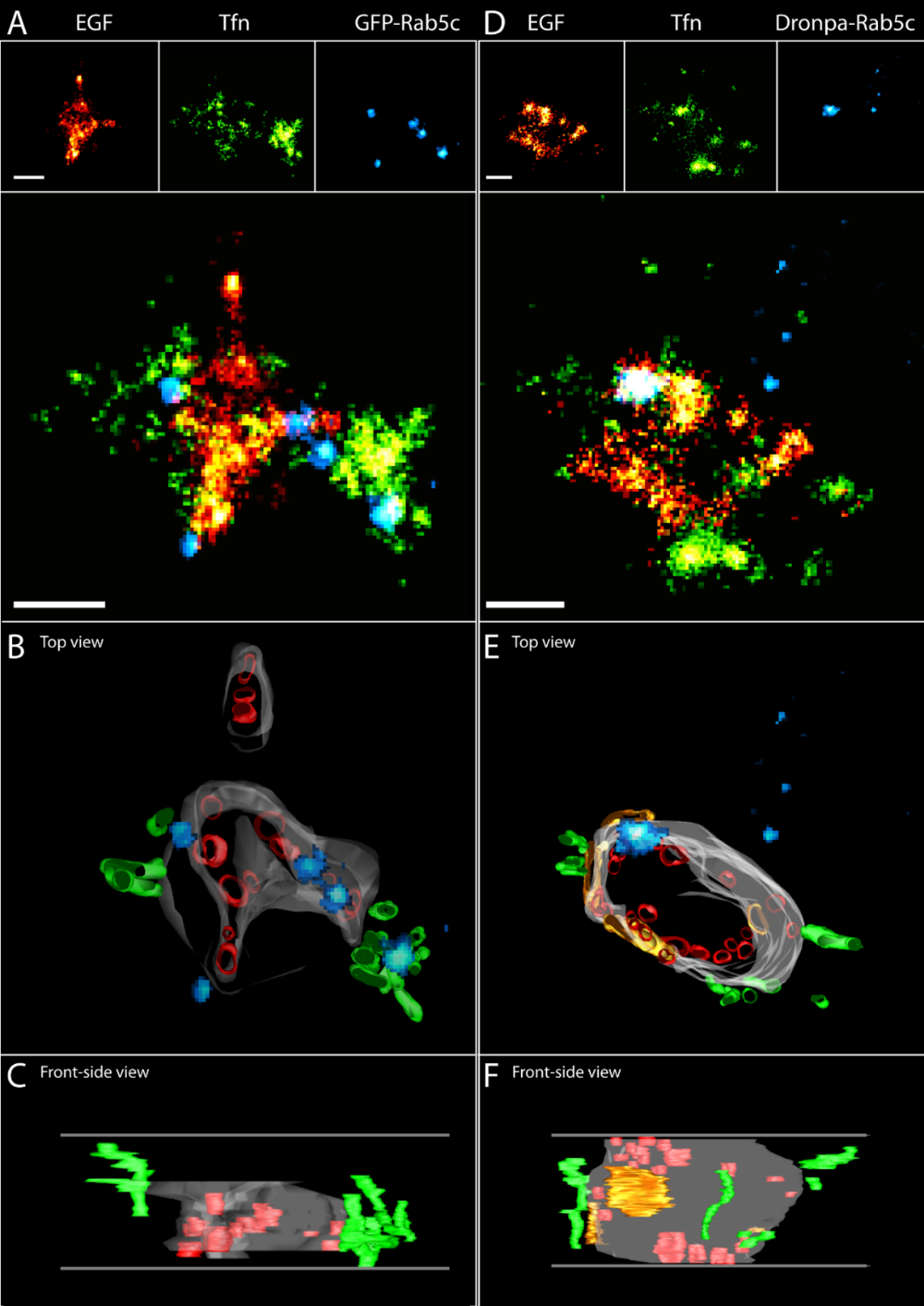

**Figure S12: Mapping of Rab5c on endosomes visualized by triple-colour superCLEM (additional examples).**

Mapping of Rab5c on endosomes visualized by triple-colour superCLEM using semi-thin Tokuyasu sections. (A,D) SMLM data for EGF-Alexa647 (red), TFn-Alexa568 (green) and GFP/Dronpa-Rab5c (blue). (B,C,E,F) Ultrastructural models of endosomes based on tomograms reconstructed from double-axis tilt series. Colours used in models represent: (grey) limiting membrane, (red) ILV, (green) recycling tubules, (orange) sorting microdomains (SµD), (blue) Rab5c. Horizontal grey lines in the bottom images (C,F) indicate top and bottom sides of reconstructed tomograms. Scalebar: 250 nm.
